# Lacking social support is associated with structural divergences in hippocampus-default network co-variation patterns

**DOI:** 10.1101/2021.08.19.456949

**Authors:** Chris Zajner, Nathan Spreng, Danilo Bzdok

## Abstract

Elaborate social interaction is a pivotal asset of the human species. The complexity of people’s social lives may constitute the dominating factor in the vibrancy of many individuals’ environment. The neural substrates linked to social cognition thus appear especially susceptible when people endure periods of social isolation: here, we zoom in on the systematic inter-relationships between two such neural substrates, the allocortical hippocampus (HC) and the neocortical default network (DN). Previous human social neuroscience studies have focused on the DN, while HC subfields have been studied in most detail in rodents and monkeys. To bring into contact these two separate research streams, we directly quantified how DN subregions are coherently co-expressed with specific HC subfields in the context of social isolation. A two-pronged decomposition of structural brain scans from ∼40,000 UK Biobank participants linked lack of social support to mostly lateral subregions in the DN patterns. This lateral DN association co-occurred with HC patterns that implicated especially subiculum, presubiculum, CA2, CA3, and dentate gyrus. Overall, the subregion divergences within spatially overlapping signatures of HC-DN co-variation followed a clear segregation divide into the left and right brain hemispheres. Separable regimes of structural HC-DN co-variation also showed distinct associations with the genetic predisposition for lacking social support at the population level.

## Introduction

Our brains are highly attuned to mediating social relationships. Yet, this unique property may make us especially vulnerable in times of social isolation. The hippocampus (HC) and the highly associative default network (DN) in particular play key roles in representing and integrating knowledge about the panoply of people’s social worlds (Spreng and Andrews-Hanna, 2015; Tavares et al., 2015). Evidence across the primate lineage demonstrates expansion of the brain towards continuously larger association cortex volumes. It has been suggested that this selective expansion of association cortex brain regions has especially coincided with the benefits of social bonding and coping with life in social groups (Dunbar, 1998; Dunbar and Shultz, 2007). Additionally, based on within-species assessments amongst humans, the size of one’s social network has been found to correlate with the structure of brain regions, including key parts of the DN (Lewis et al., 2011). This finding receives further support from monkey experiments aimed at controlling group size (Sallet et al., 2011). In this light, recently expanded regions of the association cortex appear to be related to the advanced processing capacities required for navigating social exchange with others (Schurz et al., 2021a).

However, the neocortical nodes of the DN have long been emphasized in the brain-imaging community to be integral for internally-generated or self-directed cognitive functions, as opposed to externally or environment-oriented processing (Andrews-Hanna et al., 2014). This raises the question of how the DN supports a central role in social embeddedness. Emotional connection is one fundamental aspect of social cognition. Yet, this domain is classically believed to be subserved by the limbic system, which includes the hippocampus (Schurz et al., 2021b). On the other hand, the more cognitive and abstract reasoning-based aspects of social processing may be preferentially subserved by neocortical regions of the DN, such as the medial prefrontal cortex (mPFC), inferior parietal lobule (IPL), and temporoparietal junction (TPJ). For example, these same regions are associated with general perspective-taking competence (Lewis et al., 2011), and show neural activity responses when thinking about others (Krienen et al., 2010). Thus, the relationship between the internal representation of the social world and DN regions could perhaps be a key factor behind the recent discovery that the DN is the network circuit with the strongest links to subjective social isolation (Spreng et al., 2020). Yet, we still have an incomplete understanding of how social isolation and the disparate subsystems within the higher association cortex are inter-linked with their affiliates in the allocortex.

This knowledge gap is in part due to inherent methodological challenges posed by the endeavor of studying DN regions that are particularly evolved in the human species. From a comparative perspective, progress in elucidating the DN in humans is hampered by the difficulty of confidently matching distinct DN subregions to counterparts in the brains of our monkey ancestors, or even other animals. Indeed, several neocortical areas of the human association cortex may have no definitive homolog in non-human animals (Petrides et al., 2012). For example, there are hurdles to the attempts of identifying homologous structures of the human TPJ and mPFC in the monkey brain (Mars et al., 2013; Saxe, 2006). Overall, such incongruences in comparative studies have impeded “the ability to compare experimental findings from nonhuman primates with results obtained in functional and structural neuroimaging of the human brain” (Petrides et al., 2012).

In contrast, there is an extensive knowledge base on the evolutionarily more conserved hippocampus in the allocortex (Buzsaki, 2006; Xu et al., 2020). This is due to the ready possibility for conducting direct experiments on the hippocampus of animals, which are typically infeasible in humans. For example, invasive studies in living animals using direct axonal tracing, gene expression probes, optogenetic manipulation and single-cell electrophysiological recording in the rodent and monkey hippocampus have enriched our understanding of this structure and its functionally specialized subfields. There is also accumulating knowledge of the effects of social isolation on specific subregions of the animal hippocampus (Biggio et al., 2019; Ibi et al., 2008; Kogan et al., 2000; Silva-Gomez et al., 2003). For these reasons, the neuroscientific understanding of the neocortical subregions of the DN remain more opaque than those of the hippocampus. Thus, simultaneously investigating these two functionally interacting neural systems opens a window, which can eventually allow illuminating key properties of human DN subregions through the lens of their partner hippocampus subregions.

We expected concordances in how these allocortical and neocortical neural systems are related to social isolation. This is because both circuits are implicated in serving social interaction, such as the abstract representation of other people’s purview of the world. On the one hand, the DN is well-known to be involved in both representing information about other people (Courtney and Meyer, 2020), and taking other peoples’ perspective (Frith and Frith, 2006; Jamali et al., 2021; Numssen et al., 2021; Saxe and Kanwisher, 2003). On the other hand, the hippocampus of various animal species has been shown to serve proto-forms of such functions (Danjo et al., 2018; Oliva et al., 2020; Omer et al., 2018). This idea is supported by the wide range of information domains which individual hippocampus neurons are reported to be capable of representing, as evidenced by invasive electrophysiological recordings in rodents and monkeys (Behrens et al., 2018; Bellmund et al., 2018; Eichenbaum et al., 1999). These include spatial boundaries (Lever et al., 2009), head-direction (Taube et al., 1990), goal direction (Sarel et al., 2017), sound frequency (Aronov et al., 2017), odor (Radvansky and Dombeck, 2018; Wood et al., 2000), time (MacDonald et al., 2011), and reward (Gauthier and Tank, 2018). These representations related to single-cell activity extend to social information as well. For example, in rodents and bats, individual hippocampus neurons have been shown to specifically represent the location of peers within a spatial environment (Danjo et al., 2018; Oliva et al., 2020; Omer et al., 2018).

The human hippocampus is also increasingly believed to represent discrete items of social information. For example, *in-vivo* electrophysiological experiments in epilepsy patients have shown that single hippocampus neurons in the medial temporal lobe consistently respond to pictures of the same person from diverse viewpoints (Quiroga et al., 2009; Quiroga et al., 2005; Rey et al., 2020). Additionally, a similar study of neurosurgical patients found that single hippocampus neurons tend to respond to different images of people if those images were previously judged by the patient to be similar (De Falco et al., 2016). The conjunction of these earlier studies suggests that neurons in the hippocampus of human and non-human mammals play a fundamental role in recognizing and representing specific peers. These social representations may hence be embedded within neural representations of ‘social spaces’ mediated by the entorhinal cortex.

Indeed, the entorhinal cortex ⍰ which is the main input and output structure of the hippocampus ⍰ has repeatedly been shown to code an animals’ location through an ensemble of ‘grid-cells’ (Hafting et al., 2005; Jacobs et al., 2013; Moser et al., 2008). These dedicated neurons are believed to discharge according to fields that tessellate the environment with regular hexagonal patterns. In fact, such ‘grid-cells’ have been demonstrated to map multiple different domains of information, such as space (Hafting et al., 2005; Jacobs et al., 2013; Moser et al., 2008), time (Kraus et al., 2015), and speed (Kropff et al., 2015). Some studies have also shown that the pre- and parasubiculum support such ‘grid cell’ representations (Boccara et al., 2010).

The constant representation of spatial environments by grid-cells has been further suggested as a bedrock for a long discussed role of the hippocampus ⍰ to instantiate ‘cognitive maps’, classically maps of space (O’keefe and Nadel, 1978; Tolman, 1948). Later, it was suggested that this function in spatial conceptualization has been co-opted in the primate brain to help instantiate other abstract maps of related entities (Behrens et al., 2018; Bellmund et al., 2018; Constantinescu et al., 2016). Recent evidence suggests that similar spatial maps are also represented in the orbital frontal cortex region of the DN (Wikenheiser et al., 2021). Evidence of a cognitive map of interpersonal ties has been further associated with neural activity responses of the human hippocampus. This involved both social agent ‘nodes’ and their social relationship ‘edges’ (Tavares et al., 2015). In particular, hippocampal fMRI activity could track the position of characters within a social hierarchy as indexed by ‘power’ and ‘affiliation’ (Tavares et al., 2015). Moreover, in this human fMRI experiment, individuals with better social skills showed more pronounced fMRI activity responses (Tavares et al., 2015). It is thus conceivable that advanced types of social cognition, such as perspective-taking, require accessing and binding information within an abstract social relationship ‘map’ mediated by the hippocampus. If so, we reasoned that the hippocampal subregions which play central roles in instantiating a cognitive map are potentially linked to the regular exchange in one’s wider social networks and lack thereof. The shared functions of the hippocampus and DN thus point towards principled covariation between their subcomponents.

In the past, brain-imaging studies aiming at brain parcellation have typically investigated either the default network (e.g., Schurz et al., 2014) or the hippocampus alone (e.g., Plachti et al., 2019). Although there is extensive research from animal studies on anatomically defined subregions of the hippocampus, the extension of this work to investigations on the human hippocampus are still lacking. Advances in the automatic segmentation of the hippocampus using *ex vivo* brain imaging (Iglesias et al., 2015; Wisse et al., 2017) now allow reliable assessments of micro-anatomically defined hippocampus subregions in a way that scales to the ∼40,000 UK Biobank Imaging cohort. This enables deeper analyses of the principled inter-relationships between the evolutionarily more conserved allocortical hippocampus and default network of the recently expanded neocortex. By leveraging a framework for high-dimensional decomposition at a fine-grained subregion resolution, we here uncover the structural deviations of the hippocampus-default network co-variations signatures that characterize objective social isolation. Moreover, enabled by the availability of genetic data, we investigate how the structural brain patterns are associated with the genetic predisposition to experience social isolation.

## Material and Methods

### Data resources

The UK Biobank is a prospective epidemiology resource that offers extensive behavioral and demographic assessments, medical and cognitive measures, as well as biological samples in a cohort of ∼500,000 participants recruited from across Great Britain (https://www.ukbiobank.ac.uk/). This openly accessible population dataset aims to provide brain-imaging for ∼100,000 individuals planned for completion in 2022. The present study was based on the recent data release from February/March 2020. To improve comparability and reproducibility, our study built on the uniform data preprocessing pipelines designed and carried out by FMRIB, Oxford University, UK (Alfaro-Almagro et al., 2018). Our study involved data from 38,701 participants with brain-imaging measures and expert-curated image-derived phenotypes of grey matter morphology (T1-weighted MRI) from 48% men and 52% women, aged 40-69 years when recruited (mean age 55, standard deviation [SD] 7.5 years). The present analyses were conducted under UK Biobank application number 25163. All participants provided informed consent. Further information on the consent procedure can be found elsewhere (http://biobank.ctsu.ox.ac.uk/crystal/field.cgi?id=200).

### Target phenotype for objective social isolation

To capture an objective measure of the frequency of social interactions, our UK Biobank analyses were based on answers to the question ‘How often are you able to confide in someone close to you?’(data field 2110). Our study modeled lack of social support as less than ‘daily or almost daily’ (yes or positive answer) against confiding in others more often (treated as no or negative answer).

Measures of social relationship quality represent a widely accepted and widely investigated component of social embeddedness (Cohen and Hoberman, 1983; Hawkley et al., 2005). Lack of social support is commonly viewed as an objective measure of weak social connection with other people. For examples, the Social Relationships scales are part of the NIH Toolbox (Cyranowski et al., 2013) feature dimensions of social networks, which closely resembles our measure of social support. In general, a variety of studies showed single-item measures of social isolation traits to be reliable and valid (Atroszko et al., 2015; Dollinger and Malmquist, 2009; Mashek et al., 2007). The sociodemographic differences between low and high social support individuals in the UK Biobank have been reported elsewhere (Schurz et al., 2021b).

### Brain-imaging and preprocessing procedures

Magnetic resonance imaging scanners (3T Siemens Skyra) were matched at several dedicated data collection sites with the same acquisition protocols and standard Siemens 32-channel radiofrequency receiver head coils. To protect the anonymity of the study participants, brain-imaging data were defaced and any sensitive meta-information was removed. Automated processing and quality control pipelines were deployed (Alfaro-Almagro et al., 2018; Miller et al., 2016). To improve homogeneity of the imaging data, noise was removed by means of 190 sensitivity features. This approach allowed for the reliable identification and exclusion of problematic brain scans, such as due to excessive head motion.

The structural MRI data were acquired as high-resolution T1-weighted images of brain anatomy using a 3D MPRAGE sequence at 1 mm isotropic resolution. Preprocessing included gradient distortion correction (GDC), field of view reduction using the Brain Extraction Tool (Smith 2002) and FLIRT (Jenkinson and Smith 2001; Jenkinson et al. 2002), as well as non-linear registration to MNI152 standard space at 1 mm resolution using FNIRT (Andersson et al. 2007). To avoid unnecessary interpolation, all image transformations were estimated, combined and applied by a single interpolation step. Tissue-type segmentation into cerebrospinal fluid (CSF), grey matter (GM) and white matter (WM) was applied using FAST (FMRIB’s Automated Segmentation Tool, (Zhang et al. 2001)) to generate full bias-field-corrected images. SIENAX (Smith et al. 2002), in turn, was used to derive volumetric measures normalized for head sizes.

For the default network, volume extraction was anatomically guided by the Schaefer-Yeo reference atlas (Schaefer et al., 2017). Among the total of 400 parcels, 91 subregion definitions are provided as belonging to the default network among the 7 canonical networks. For the hippocampus, 38 volume measures were extracted using the automatic Freesurfer sub-segmentation (Iglesias et al., 2015). The allocortical volumetric segmentation draws on a probabilistic hippocampus atlas with ultra-high resolution at ∼0.1mm isotropic. This tool from the Freesurfer 7.0 suite gives special attention to surrounding anatomical structures to refine the hippocampus subregion segmentation in each participant.

As a preliminary data-cleaning step, building on previous UK Biobank research (Schurz et al., 2021b; Spreng et al., 2020), inter-individual variation in brain region volumes that could be explained by nuisance variables of no interest were regressed out: body mass index, head size, head motion during task-related brain scans, head motion during task-unrelated brain scans, head position and receiver coil in the scanner (x, y, and z), position of scanner table, as well as the data acquisition site, in addition to age, age^2^, sex, sex*age, and sex*age^2^. The cleaned volumetric measures from the 91 DN subregions in the neocortex and the 38 HC subregions in the allocortex served as the basis for all subsequent analysis steps.

### Analysis of co-variation between hippocampus subregions and default-network subregions

As the keystone of the analytical workflow, we sought dominant regimes of structural correspondence – signatures or “modes” of population co-variation that provide insights into how *structural variation among the segregated HC* can explain *structural variation among the segregated DN*. Canonical correlation analysis (CCA) was a natural choice of method to interrogate such multivariate inter-relations between two high-dimensional variable sets (Bzdok et al., 2019; Wang et al., 2020; Witten et al., 2009).

A first variable set *X* was constructed from the DN subregion volumes (number of participants × 91 DN parcels matrix). A second variable set *Y* was constructed from the HC subregion volumes (number of participants × 38 HC parcels matrix):

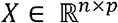

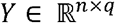

where *n* denotes the number of observations or participants, *p* is the number of DN subregions, and *q* is the number of HC subregions. Each column of the two data matrices was z-scored to zero mean (i.e., centering) and unit variance (i.e., rescaling). CCA addresses the problem of maximizing the linear correlation between low-rank projections from the two variable sets or data matrices. The two sets of linear combinations of the original variables are obtained by optimizing the following target function:

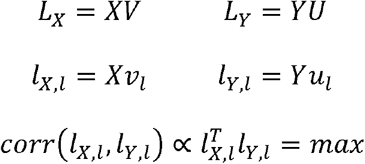

where *V* and *U* denote the respective contributions of *X* and *Y*, *L_X_* and *L_X_* denote the respective latent ‘modes’ expression of joint variation (i.e., canonical variates) based on patterns derived from *X* and patterns derived from *Y*, *l_X,l_* is the *l*th column of *L_X_*, and *l_Y,l_* is the *l*th column of *L_Y_*. The goal of our CCA application was to find pairs of latent vectors *l_X,l_* and *l_Y,l_* with maximal correlation in the derived latent embedding. In an iterative process, the data matrices *X* and *Y* were decomposed into *L* components, where *L* denotes the number of modes given the model specification. In other words, CCA involves finding the canonical vectors *u* and *v* that maximize the (symmetric) relationship between a linear combination of DN volume expressions (*X*) and a linear combination of HC volume expressions (*Y*). CCA thus identifies the two concomitant projections *Xv_l_* and *Yu_l_*. These yielded the optimized co-occurrence between combined subregion variation inside the segregated DN and combined subregion variation inside the segregated HC across participants.

In other words, each estimated cross-correlation signature identified a constellation of within-DN volumetric variation and a constellation of within-HC volumetric variation that go hand-in-hand with each other. The set of *k* orthogonal modes of population co-variation are mutually uncorrelated by construction (Wang et al., 2020). They are also naturally ordered from the most important to the least important HC-DN co-variation mode based on the amount of variance explained between the allocortical and neocortical variable sets. The first and strongest mode therefore explained the largest fraction of joint variation between combinations of HC subregions and combinations of DN subregions. Each ensuing cross-correlation signature captured a fraction of structural variation that is not explained by one of the *k* - 1 other modes. The variable sets were entered into CCA after a confound-removal procedure based on previous UK Biobank research (cf. above).

### Group difference analysis

For each of the derived population modes of HC-DN co-variation, we then performed a rigorous group contrast analysis for social isolation. We aimed to identify which anatomical subregions show statistically defensible deviation in the socially isolated group compared to the control group. For the low social support trait, we carried out a principled test for whether the solution vector obtained from CCA (i.e., canonical vectors, cf. above) in the socially isolated group is systematically different from the solution vector in the control group.

More specifically, following previous UK Biobank research (Schurz et al., 2021b), we carried out a bootstrap difference test of the CCA solution from the socially isolated group vs. socially well-connected group (Efron and Tibshirani, 1994). In 100 bootstrap iterations, we randomly pulled participant samples with replacement to build an alternative dataset (with the same sample size) that we could have gotten. We subsequently performed CCA in parallel by fitting one separate model to each of the two groups. In each resampling iteration this approach thus carried out a separate estimation of the doubly multivariate correspondence between HC subregions and DN subregions in each particular group. The two distinct CCA solutions from each iteration were then matched mode-by-mode regarding sign invariance and mode order. Canonical vectors of a given mode that carried opposite signs were aligned by multiplying one with −1. The order of the CCA modes was aligned based on pairwise Pearson’s correlation coefficient between the canonical vectors from each estimated CCA model. After mode matching, we directly estimated the resample-to-resample effects by elementwise subtraction of the corresponding canonical vectors of a given mode *k* between the two groups. We finally recorded these difference estimates for each vector entry (each corresponding to the degree of deviation in one particular anatomical subregion). The subregion-wise differences were ultimately aggregated across the 100 bootstrap datasets to obtain a non-parametric distribution of group contrast estimates.

We thus propagated the variability attributable to participant sampling into the computed uncertainty estimates of group differences in the UK Biobank population cohort. Statistically relevant alteration of anatomical subregions in social isolation were determined by whether the two-sided confidence interval included zero or not according to the 10/90% bootstrap-derived distribution of difference estimates (Schurz et al., 2021b). Remaining faithful to our multivariate analytical strategy and research question, this non-parametric approach directly quantified the statistical uncertainty of how social isolation is manifested in specific subregions of the HC-DN axis.

### Analysis of how individual expressions of HC-DN co-variation are linked to the genetic predisposition for social isolation

Polygenic risk scores (PRS) are a genome-wide analysis technique that has been shown to successfully quantify the genetic predisposition of individuals for a variety of phenotypes. The approach has become especially potent for complex phenotypes that implicate tens of thousands of common-variant single nucleotide polymorphisms (SNPs) with individually small effect sizes, such as major psychiatric diseases (Elliott et al., 2020; HLA-C, 2009; Inouye et al., 2018; Khera et al., 2018; Kuchenbaecker et al., 2017). PRS have also come to be a sharp tool for heritability analyses due to the advent of large population datasets (e.g., the UK Biobank) (Choi et al., 2020; Wray et al., 2021). Such data resources have allowed the investigation of the relationship between SNP-based genetic variation and inter-individual differences in a particular phenotype, which includes neuroimaging-derived phenotypes (Elliott et al., 2018; Smith et al., 2021). For the purpose of the present study, we have constructed a PRS model for the low social support trait. The subject-specific risk scores were then regressed onto our expressions of HC-DN modes (i.e., canonical variates). Our integrative imaging-genetics approach aimed to disentangle which mode expressions have reliable links to the genetic vulnerability to low social support trait as attributable to thousands of genetic variants.

As is common for PRS analysis workflows, summary statistics from previously conducted genome-wide association studies (GWAS) on our target phenotypes were used as the backdrop to determine how several hundred thousand SNPs are associated with the low social support trait. The summary statistics for low social support were obtained from a GWAS that was conducted as part of the Canadian Longitudinal Study on Aging. Quality control was implemented by excluding SNPs with a minor allele frequency of less than 1%, as well as excluding SNPs with a difference between reported sex and that indicated by their sex chromosomes, and removing overlapping samples.

The quality-controlled summary statistics were used as starting point for the PRS model that was built and applied using the PRSice framework (http://www.prsice.info). This software tool uses the available collection of effect sizes of candidate SNPs to form single-subject predictions of genetic predisposition for a phenotype of interest. More specifically, this tool determined the optimal PRS model based on the UK Biobank participants (training data, n=253,295) of European ancestry who did not provide any brain-imaging data (at the time of study). This model training step involved automated adjustments, such as identifying ideal clumping and pruning choices, to select the thresholds that decide which SNPs are included in the PRS model. Subsequently, once optimized, the final PRS model was then used to predict the genetic predisposition for each of 23,423 UK Biobank participants of European ancestry *with* brain-imaging data (test data). This PRS model consisted in pooling across additive effects of weighted SNPs, whereby the weighted sum of the participants’ genotypes was computed as follows:

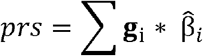

where g_i_ denotes an individual’s genotypes at SNP *i* (values 0, 1, or 2), 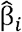 is the obtained point estimates of the per-allele effect sizes at SNP *i*, and *j* is a particular individual (Choi et al., 2020).

Finally, Bayesian linear regression was used to regress the subject-specific predictions of genetic liability for the low social support trait onto the participant expressions of HC-DN co-variation modes. To this end, the individuals in the top 5% predictions (i.e., highest PRS estimates) and the individuals in the bottom 5% predictions (i.e., lowest PRS estimates) were considered as a binary outcome in a Bayesian logistic regression model with participant-wise mode expressions serving as input variables (Fan et al., 2019; Lecarpentier et al., 2017; Meisner et al., 2020). In this multiple regression setup, PRS for low social support was regressed against each of the 25 canonical variates (linearly uncorrelated by construction) on the HC side for every individual. An analogous multiple regression model was estimated for the (uncorrelated) 25 canonical variates from the DN side. The fully Bayesian model specification for these regression analyses was as follows:

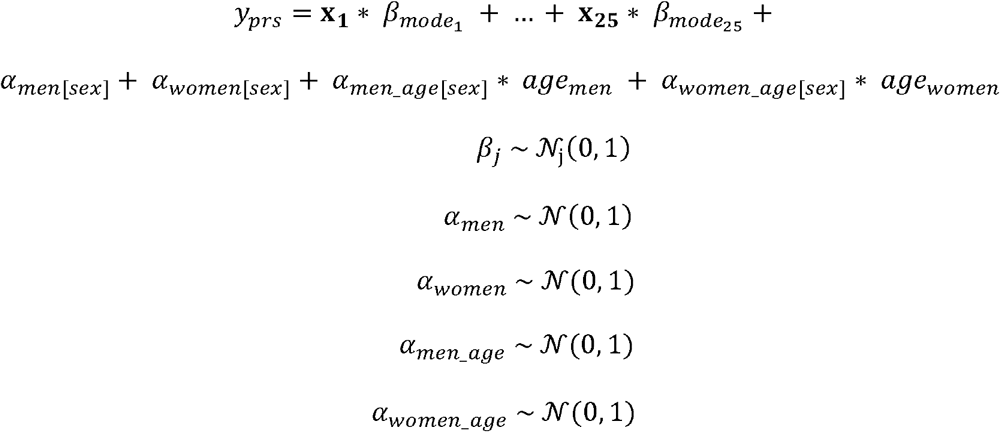

where *β_j_* denotes the slopes for the subject-specific 25 mode expressions as canonical variates x_j_, and *y_prs_* denotes the PRS estimates of each participant. Potential confounding influences were acknowledged by the nuisance variables α, which accounted for variation that could be explained by sex and (z-scored) age. Once the Bayesian model solution was approximated using Markov chain Monte Carlo sampling, it yielded fully probabilistically specified posterior parameter distributions for each *β* coefficient corresponding to one of the signatures of allocortical-neocortical co-variation (cf. above). The association with trait heritability of a mode expression was then determined based on how robustly their corresponding model coefficients deviated from 0 (e.g., >95% of model coefficient posterior probability excluded a value of 0).

## Results

### Structural co-variation between hippocampus and default network at the subregion level

We explored the principal signatures of structural co-variation between the full set of 38 hippocampal subregions and the full set of 91 DN subregions. The concurrent patterns within subregion variation among the hippocampus and those within the DN were computed using a doubly multivariate learning algorithm. In so doing, we achieved a co-decomposition of hippocampal subregion volumes and DN subregion volumes. Each of the top 25 modes of co-variation was characterized based on how much of joint variance a particular signature explained: with the most explanatory signature (mode 1) achieving a canonical correlation of rho = 0.51 (measured as Pearson’s correlation coefficient) (see Supplementary Table 1). The second most explanatory signature (mode 2) achieved a canonical correlation of rho = 0.42, the third signature rho = 0.39, the fourth signature rho = 0.31, the fifth signature rho = 0.27, and the sixth rho = 0.23; through to the twenty-fifth signature which had rho = 0.06 (see Supplementary Table 1 for full list). This analysis thus established the scaffold for all subsequent analyses that delineates how multiple complementary hippocampal patterns co-vary hand-in-hand with DN patterns across individuals.

### Differences in the hippocampus-default network co-variation in social isolation

Based on the identified population signatures of HC-DN co-variation, we investigated the neurobiological manifestations of social isolation in our UK Biobank sample. This was accomplished by examining robust subregion-level divergences in how hippocampal patterns are co-expressed with DN patterns that characterize groups of socially isolated vs. control participants (i.e., low vs. high social support). To this end, we analyzed objective social isolation by a rigorous group difference analysis between the structural patterns of co-variation in the low social support and high social support groups. This approach revealed the precise subregions contributing to the structural HC-DN co-variation that systematically diverged between the two groups, for each mode of the CCA.

We uncovered a multitude of modes with systematic group differences in either specific HC and/or DN subregions. We also found modes with no significant structural divergences in any HC or DN subregion. From here on, a subregion which was observed to have a robustly different weighting within a modes canonical vector, between low and high social support groups, is termed a ‘hit’ (i.e., an observed structural divergence in individuals with low social support). Across all 25 examined modes, contrasting low vs. high social support, we identified hits in 32 HC subregions and 50 hits in DN subregions. Most of these hits occurred in earlier modes, with all the hits occurring between modes 1-7. Just in the first three modes, we found 24 HC subregion hits (70.6% of the HC total). In the first mode alone, we also observed 26 DN hits (52% of the DN total). The largest number of HC hits were observed in presubiculum (5 hits), subiculum (5), CA2/3 head (4), and GC-DG-ML (4). Regarding the parallel DN divergences, we found the largest number of hits in lateral cortical subregions (78% of the total), such as the middle temporal gyrus (MTG), temporal pole, and IPL, with less in midline subregions. There was a total number of hits in 17 temporal (34%), 17 prefrontal (34%), 11 parietal (22%), and 5 posterior cingulate (10%) subregions.

The divergences between the low versus high social support groups for mode 1 (see Figure 1) revealed a rough synopsis of the hit locations for the totality of the observed modes. In the dominant mode, we observed hits in bilateral CA2/3 head and body, bilateral CA4 head, left CA4 body, bilateral HATA, left presubiculum head, bilateral subiculum body, and bilateral GC-DG-ML head and body, with 26 hits in DN subregions (8 temporal, 7 parietal, 10 prefrontal, 1 posterior cingulate). Additionally, the HC subregion hits with the strongest weights included left CA2/3 body and bilateral GC-DG-ML. On the flipside of our model, the DN subregion hits with the strongest weights were bilateral prefrontal cortex, left MTG, and left IPL.

In the second most explanatory signature of HC-DN co-variation (i.e., mode 2), we identified 7 HC hits ⍰ bilateral tail, bilateral parasubiculum, bilateral presubiculum head, and left subiculum body ⍰ and no hits in DN subregions (see Supplementary Figure 1). In mode 3, we observed 2 HC hits located in left CA2/3 head, and right subiculum head. We also observed 7 DN hits, located in the ventromedial prefrontal cortex (vmPFC), dorsolateral prefrontal cortex (dlPFC), and TPJ ⍰ all of which were in the left hemisphere (see Figure 2). In mode 4, we observed 3 HC hits located in left presubiculum head, left fimbria, and left HATA. All these HC hits for mode 4 were in the left hemisphere. Conversely, we observed 4 DN hits exclusively in the right hemisphere (see Figure 3). These DN hits emerged in the temporal pole, retrosplenial cortex (RSC), anterior cingulate cortex, and dlPFC. In mode 5 we observed 4 HC hits located only in the right hemisphere. These hits included the fimbria, presubiculum head, subiculum head, and molecular layer head. We also noted one hit in the DN: right posterior cingulate cortex (PCC) (see Supplementary Figure 2). In mode 6, we observed no HC hits and 12 DN hits all in the left hemisphere (see Figure 4). These hits included 8 in the lateral temporal lobe, 2 in the IPL, and 2 in the RSC. In mode 7, we observed a hit in the left parasubiculum and no DN hits (see Supplementary Figure 3). Beyond mode 7, we did not observe hits in any of the other 25 examined modes. Thus, the more dominant signatures of HC-DN covariation showed stronger relationships to a lack of social support than less dominant signatures. We present here the modes with the greatest number of subregion divergences in both the HC and DN (see Figures 1–4). These collective results show that a group contrast analysis of low vs. high social support individuals revealed specific subregion divergences within spatially overlapping signatures of HC-DN co-variation.

Overall, we noted a recurring theme of certain subregions with numerous hits in the group analysis of social support. For the hippocampus, this included the presubiculum, subiculum, CA2/3, and GC-DG-ML. For the DN, especially lateral subregions ⍰ such as the dlPFC, IPL and MTG ⍰ tended to diverge between low vs. high social support groups. We also noted that the divergences observed for low social support occurred in structural patterns within each mode, as most hits were chiefly seen on only one brain hemisphere. For example, in mode 6 there was a cluster of hits in lateral cortical regions, but only in the left hemisphere of the brain (see Figure 4). Analogously, in mode 3, we observed 7 DN hits, yet all in the left hemisphere. Thus, low social support was primarily concomitant with divergences in left lateral DN subregions, as well as the presubiculum, subiculum, CA2/3 and the GC-DG-ML of the hippocampus.

### Genetic predisposition for social isolation

We finally sought to interrogate whether the uncovered expressions of HC-DN co-variation featured systemic relationships with the participants’ liability for low social support (cf. methods). For this purpose, we computed polygenic risk score predictions for the innate risk of low social support for our UK Biobank participants. We observed that there was a relevant relationship between low social support PRS and participant expressions (i.e., canonical variates) of modes 1 and 16 for the HC, and modes 3 and 11 for the DN (see Figure 5). The HC-DN modes with robust relationships were thus the relatively more dominant signatures amongst the 25 examined. The HC and DN patterns with relationships to PRS for low social support were also in modes in which we observed structural divergences of specific subregions in individuals with low social support. For example, mode 1 on the hippocampus side showed 15 subregion hits (see Figure 1), and mode 3 on the DN side showed 7 subregion hits (see Figure 2). Overall, we found that specific mode expressions in low social support individuals had robust ties to purely genetic factors, as captured by genome-wide risk predictions across tens of thousands of genetic markers.

## Discussion

We have tailored an analytical framework to examine how the anatomical substrates of HC-DN correspondence systematically deviate in individuals with a lack of regular social exchange with close others. Our approach allows extending the emerging interpretations of distinct subsystems within the DN by establishing robust cross-links with anatomically defined hippocampus subfields at the population level. We work towards this goal by directly estimating principled co-variation signatures which delineate how hippocampus subregion volumes are co-expressed with DN subregion volumes in ∼40,000 participants from the UK Biobank imaging-genetics cohort. In so doing, our study aimed to deepen the understanding of the human DN by anchoring its variation in corresponding substrates of the allocortical HC ⍰ a structure that has been implicated in mnemonic and associative processes in non-human animals, and which likely underpins human social navigation.

Past literature supports the notion that the frequency with which an individual interacts with other people resonates with the structural characteristics of the hippocampus and DN. In fact, DN subregions are implicated in representations of oneself and social others (Laurita et al., 2020; Mars et al., 2012). Likewise, the human hippocampus has been proposed to represent information about individual people (Quiroga et al., 2005), in addition to its classic functional interpretations of representing places and retrieving memories (O’Keefe and Dostrovsky, 1971; Scoville and Milner, 1957). Previous authors have even argued that the hippocampus may mediate an abstract social cognitive map for relationships between people, in the form of ‘vector angle’ representations (Tavares et al., 2015). In light of this idea, we expected individuals with lower frequency of social interactions, or low social support, to exhibit differences in specific signatures of HC-DN co-variation, compared to individuals with higher levels of social connection and support.

In accord with our expectations, we identified a set of subregional divergences that were characteristic for individuals with low social support: presubiculum, subiculum, CA2/3, GC-DG-ML and CA4 of the hippocampus. In particular, the expressions of structural co-variation across all modes revealed a total of 5 presubiculum hits, 5 subiculum, 4 CA2/3, 4 GC-DG-ML, and 3 in CA4 for individuals who lack social interaction in everyday life. Effects in these 5 subregions alone constituted as much as 66% of the total observed subregion divergences. Experimental animal models of subregional specialization within the hippocampus suggest that this may reflect differences in HC subfields which represent discrete elements of social information (Watarai et al., 2021). Such divergences would be in accord with the relationship between an individual’s objective level of social embeddedness and the richness of their neural representations of social information (Chadwick et al., 2014; Li et al., 2013; Watarai et al., 2021).

More specifically, the CA2 subregion ⍰ which here yielded 4 total hits across modes ⍰ has been shown to be integral for successful peer recognition in rodents (Alexander et al., 2016; Hitti and Siegelbaum, 2014; Leroy et al., 2018; Meira et al., 2018; Stevenson and Caldwell, 2014). For example, in mice, functional inhibition of dorsal CA2 cells through viral neurotoxin injection has been found to reduce interaction time with familiar peers (Hitti and Siegelbaum, 2014). Yet, this direct intervention on hippocampus tissue did not appear to affect sociability or other HC-dependent behaviors (Hitti and Siegelbaum, 2014). Similarly, it has been reported that mice showed an altered rate of place cell remapping in CA2 after the introduction of a peer into their environment. Conversely, exposure to a novel toy, as an enrichment of the physical environment, did not produce such effects (Alexander et al., 2016).

Individual cells in the dorsal CA1 have been found to support a similar function as they have been reported to reliably represent the spatial location of peers (Danjo et al., 2018; Omer et al., 2018), however CA2/3 cells were not recorded. Evidence that CA2 neurons especially encode information of newly encountered peers and possess place fields sensitive to social information has also recently been reported (Oliva et al., 2020). Numerous studies across species have additionally suggested that neurogenesis – a process localized to the granule cell layer of the dentate gyrus ⍰ is particularly sensitive to the experience of social isolation (Biggio et al., 2019; Dranovsky et al., 2011; Ibi et al., 2008; Stranahan et al., 2006) and social stress (Anacker et al., 2018; Gould et al., 1997; Gould et al., 1998). Hippocampal neurons thus appear to play a committed role in instantiating and integrating representations of social agents. Our results are consistent with this idea, as we find preferential hits in CA2/3 and DG, indicating systematic differences in the neural representations of social information. Neural activity and structure of the human DG and CA2/3 may thus reverberate with the richness of the environment more broadly, including social agents and relationships between them.

Similarly, CA2/3 and the DG have been discussed to be instrumental in disambiguating sensory inputs from similar stimuli (Kesner and Rolls, 2015; Knierim and Neunuebel, 2016). This is a function which has been termed ‘pattern separation’, and has been established at a single-cell level in the rodent hippocampus (Gilbert et al., 2001; Leutgeb et al., 2007; Leutgeb et al., 2004) and neuroimaging in humans (Baker et al., 2016; Bakker et al., 2008; Berron et al., 2016; McHugh et al., 2007). An emerging view is that pattern separation is a key function altered in Alzheimer’s disease (AD) (Ally et al., 2013; Parizkova et al., 2020; Wesnes et al., 2014), while AD has analogously been linked with social isolation (Marioni et al., 2015; Penninkilampi et al., 2018; Shen et al., 2021; Xiang et al., 2021). For example, in a mouse model of AD, the investigators reported reduced Aβ synaptotoxicity in the hippocampus (Li et al., 2013), and reduced cognitive impairment (Jankowsky et al., 2005) with environmental enrichment ⍰ an important source of which in humans is regular social contact. Recently, invasive cell-recording experiments also showed that AD-model mice have a selective impairment in pattern separation as subserved by the DG and CA3 (Rechnitz et al., 2021). Overall, the neurobiological underpinnings for representing a rich environment, perhaps best constituted by mapping social networks with their ties among people, may be intimately linked with AD and its accompanying cognitive consequences. Our study speaks to an association of divergences in the DG and CA2/3 with objective social isolation. This population-level insight highlights these allocortical subregions and their neocortical affiliates as important targets for future investigations on pattern separation-dependent functions and AD.

Of course, neural representation of social information and social processing are not the sole provenance of HC subregions, but rather involve dynamic functional coupling with the neocortex to support social labeling and the construction of more complex models of the social environment (Mars et al., 2021). Our analytical approach was designed to directly investigate this aspect of allocortical-neocortical correspondence in the service of social functioning. Our results underscore structural divergences in left lateral temporal and lateral parietal subregions of the highly associative DN in individuals who lack regular social exchange with others ⍰ neocortical divergences that our approach revealed to be concomitant with the allocortical divergences in CA2/3 and DG. We speculate that these conjoint divergences in left lateral temporal and parietal subregions ⍰ which are implicated in social semantics and spatial processing respectively (Alcalá-López et al., 2018; Hartwigsen et al., 2021; Mars et al., 2012) ⍰ are linked to the previously reported functional roles served by CA2/3 and DG in social cognition. This includes roles such as binding information about social relationships within a social cognitive map.

Further, we observed subregion divergences in individuals with objective social isolation to occur in almost exclusively lateralized patterns. These were consistent with left lateralized semantic processes (Binder et al., 2009; Liu et al., 2009; Numssen et al., 2021). Indeed, the vast majority of our observed DN hits were in left-hemispheric subregions in modes 1, 3, and 6 (76% of the total), consistent with the idea that sociality is fundamentally dependent on semantics as well as conceptual and symbolic (e.g. language) processing (Dunbar and Shultz, 2007; Frith and Frith, 2010; Lord, 2013). For example, much of the everyday stimulation in people’s lives is driven by social information (Mar et al., 2012; Mesoudi et al., 2006). Low access to social exchanges with other people may therefore be appropriately viewed as a condition for an overall stimulation-poor environment. Indeed, several past studies have made evident that our highlighted DN subregions are neurobiological substrates of environmental vibrancy. As some of many examples, volume and density of temporal lobe gray matter ⍰ especially the middle temporal gyrus and superior temporal sulcus ⍰ have been shown to track social network size, both as measured by online interactions in humans (Kanai et al., 2012), as well as in real-world social groups of monkeys (Sallet et al., 2011).

The anterior portion of the human hippocampus has also been proposed to be a locus of both semantic (Schacter and Wagner, 1999; Strange et al., 2014) and social (Morton et al., 2021; Vogel et al., 2020) processing. In line with this notion, we identified 21 total hits in the head portion of the hippocampus (66%), and only 11 (34%) in its body portion across all modes. The frequency of meaningful encounters with close others thus appears to be associated with structural covariance patterns of the anterior hippocampus. When considered in conjunction with the preferential structural divergences of left-lateralized DN subregions, these two broad patterns reconcile previous reports of sub-specializations within each of these two brain systems (Andrews-Hanna et al., 2010; Fanselow and Dong, 2010). In sum, social cognition draws upon social concept representation and abstract cognitive maps among other processes, which are revealed to be subserved by the concord of lateral subregions of the DN with CA2/3, DG, and the broader anterior hippocampus.

The DN has also been reliably dissociated into subsystems (Andrews-Hanna et al., 2010; Braga and Buckner, 2017; Braga et al., 2020; Braga et al., 2019; DiNicola et al., 2020). Our population neuroscience results confirm and detail this notion by uncovering distinct signatures of structural deviation in social isolation that aligns well with one of these subsystems. More specifically, our results implicate structural deviations in lateral DN subregions in objective social isolation, while past studies have highlighted together the lateral temporal cortex, TPJ, dmPFC, and temporal pole. Nonetheless, our study goes beyond previous attempts to sub-divide the DN, as we extend the interpretational themes of each DN subregional pattern by appreciating, and explicitly modeling, its consistent structural relationships with dedicated hippocampal subregions. In particular, the hippocampal patterns robustly linked with lateral temporal and parietal subregions tended to highlight CA2/3, GC-DG-ML, and CA4. Overall, these observations reinforce the idea of biologically coherent cross-dependencies that exist between these subregions of the DN and HC.

Since there appears to be an inherent relationship between objective social isolation and the HC-DN correspondence highlighted in our results, we suspected there may be a heritable contribution to these characteristics. To this end, we conducted the first ⍰ to our knowledge ⍰ polygenic risk score analysis for low social support. This analysis allowed us to investigate the genetic contributions to low social support that are due to individually small-effect size single nucleotide polymorphisms. Overall, we found that participant-specific expressions of HC-DN signatures show differing links to the heritable components of low social support. This is in accord with some previous research showing that social isolation has consistent, yet small genetic underpinnings (Spithoven et al., 2019). However, the majority of past investigations of the genetics of social isolation have focused on the subjective feeling of social connection (i.e., loneliness) (Abdellaoui et al., 2019; Boomsma et al., 2005; Day et al., 2018; Gao et al., 2017).

Yet, an underlying genetic contribution to objective levels of social support has been supported by recent genome-wide correlation analyses. For example, one study has demonstrated that our UK Biobank social support trait significantly shared genetic factors with 52 different demographic, lifestyle, and disease phenotypes (Schurz et al., 2021b). Such recent population neuroscience research indicates that the genetic determinants underlying social isolation are probably quite polygenic and involve complex gene-environment interactions. In our present study, we extended these insights by performing a PRS analysis for low social support. We furthered this by elucidating the relationship between a participant-specific predisposition for lacking social support and the brain expression of each HC-DN co-variation signature. Importantly, we found that only selected signatures of HC-DN co-variation showed a relevant relationship with genetic liability for low social support. These precise modes additionally showed numerous subregion divergences in our group difference analysis, which roughly overlapped the subregion divergences observed in low social support (i.e., modes 1 and 3). For example, expression of mode 1 on the HC side was linked to PRS for low social support and highlighted the CA2/3 and DG subregions. Participant-specific expressions of mode 3 on the DN side were also linked with PRS for low social support and showed 7 DN subregions with structural divergences, all in the left hemisphere. The genetic predisposition for low social support is thus manifest in brain networks that follow a pronounced left-right divide, reminiscent of the left-lateralized nature of semantic networks that subserve human-defining cognition (Binder et al., 2009; Liu et al., 2009; Numssen et al., 2021).

In conclusion, enabled by the breadth and depth of the UK Biobank resource, our analytical approach showed that an objective measure of social connection has robust structural concomitants in human HC and DN subregions. These allocortical-neocortical structural deviations included presubiculum, subiculum, CA2/3, and DG subregions of the hippocampus, and lateral subregions of the DN. Our framework extends understanding of the functional sub-systems of the DN and their potential preferential relationship with dedicated HC subfields. We also found that distinct signatures of HC-DN structural co-variation have robust relationships with individuals’ genetic predisposition for social disconnection. Future investigations into the genetic determinants and environmental influences on social isolation represent important research directions to further elucidate the structurally heterogenous qualities of the socially isolated brain.

## Acknowledgements

We are grateful to Adrien Peyrache and David Redish for useful comments on a previous version of the manuscript.

This project has been made possible by the Brain Canada Foundation, through the Canada Brain Research Fund, as well as by NIH grant R01AG068563A and the Canadian Institutes of Health Research (CIHR). DB was also supported by the Healthy Brains Healthy Lives initiative (Canada First Research Excellence fund), by the CIFAR Artificial Intelligence Chairs program (Canada Institute for Advanced Research), as well as Research Award and Teaching Award by Google.

## Author contributions

DB led data analysis. DB conceived and designed research; CZ and DB performed experiments; CZ and DB analyzed and interpreted the results; CZ and DB prepared figures; CZ and DB drafted manuscript; CZ, RNS, and DB edited and revised manuscript; CZ, RNS, and DB approved final version of manuscript

---

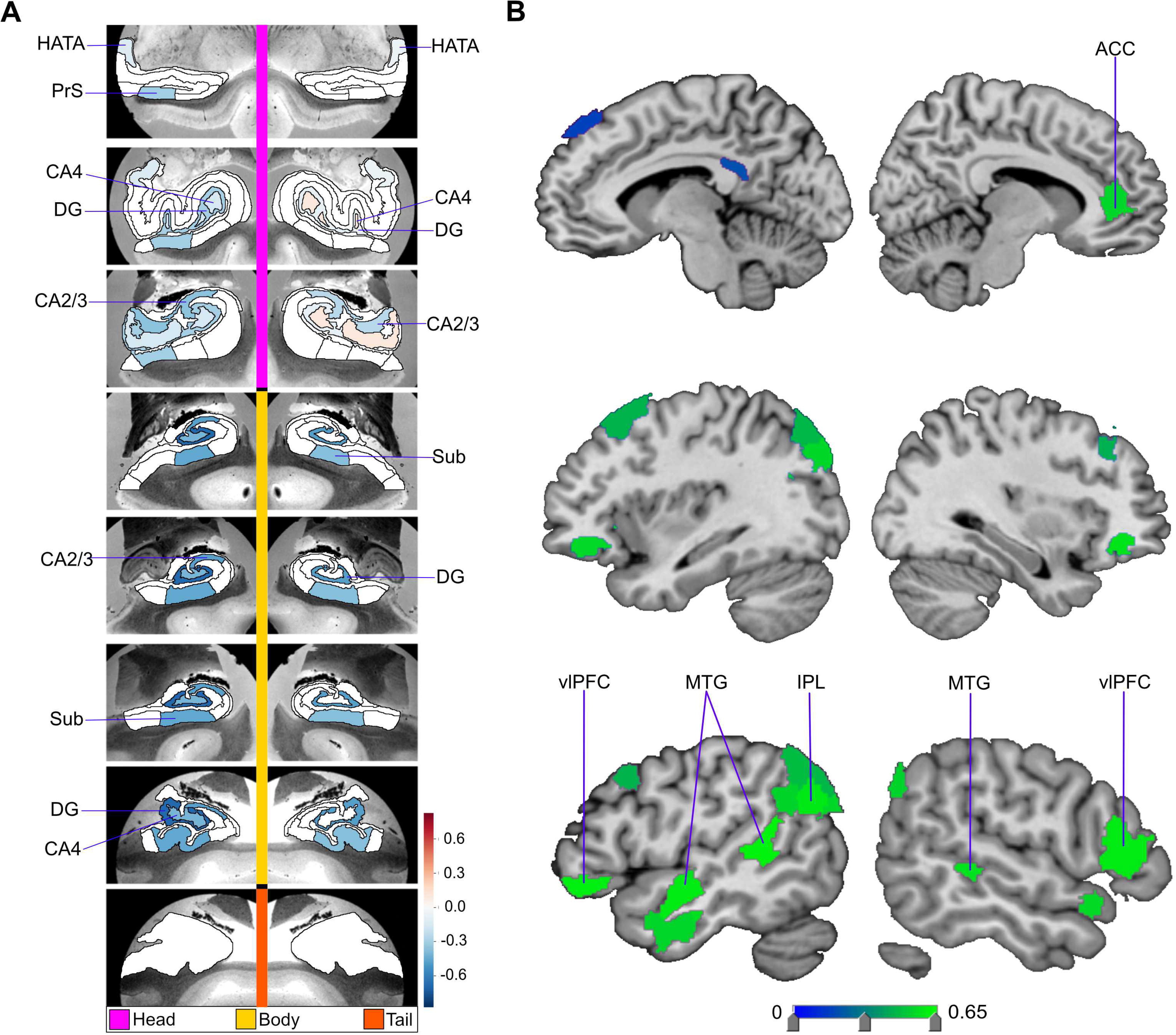

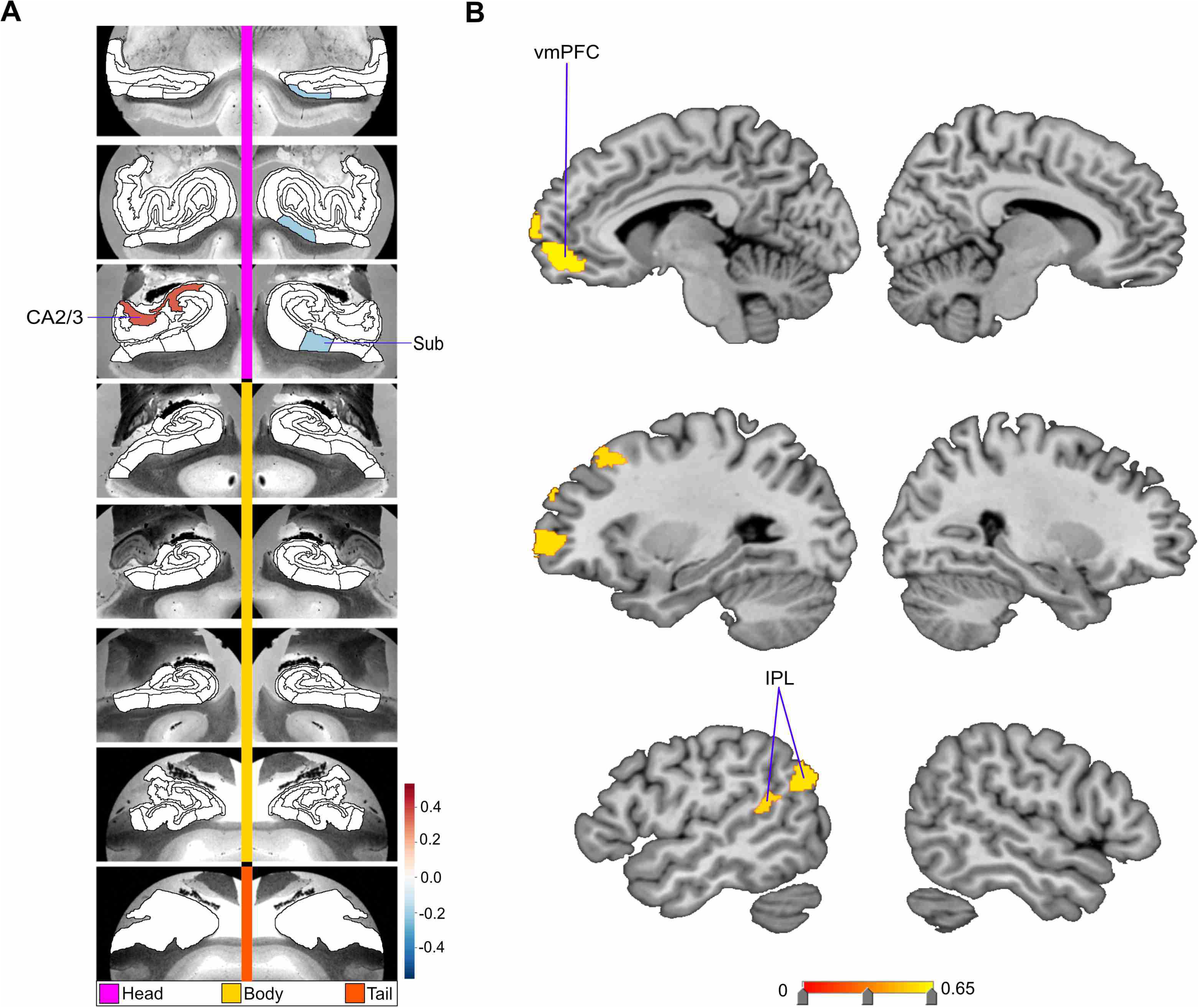

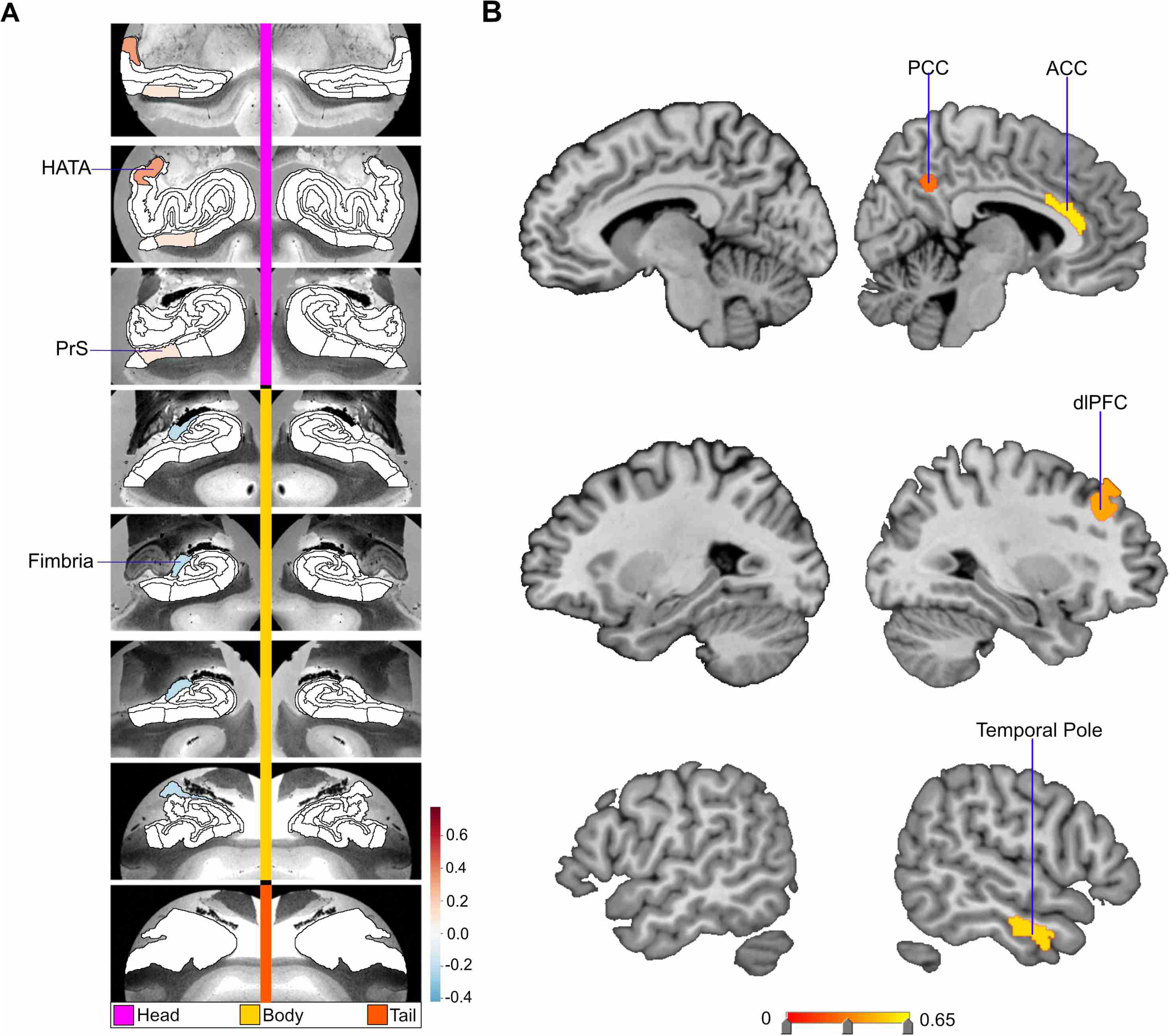

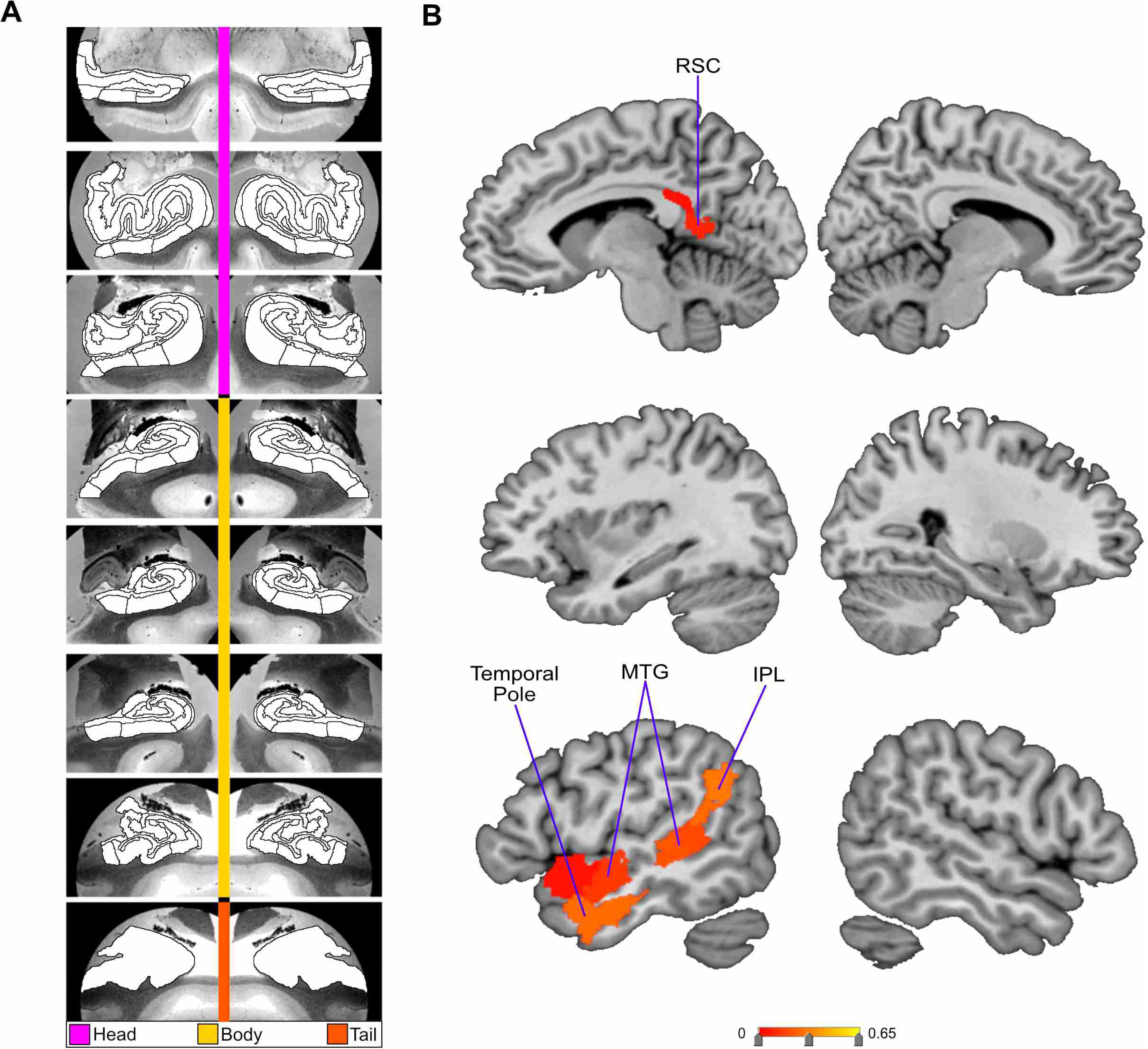

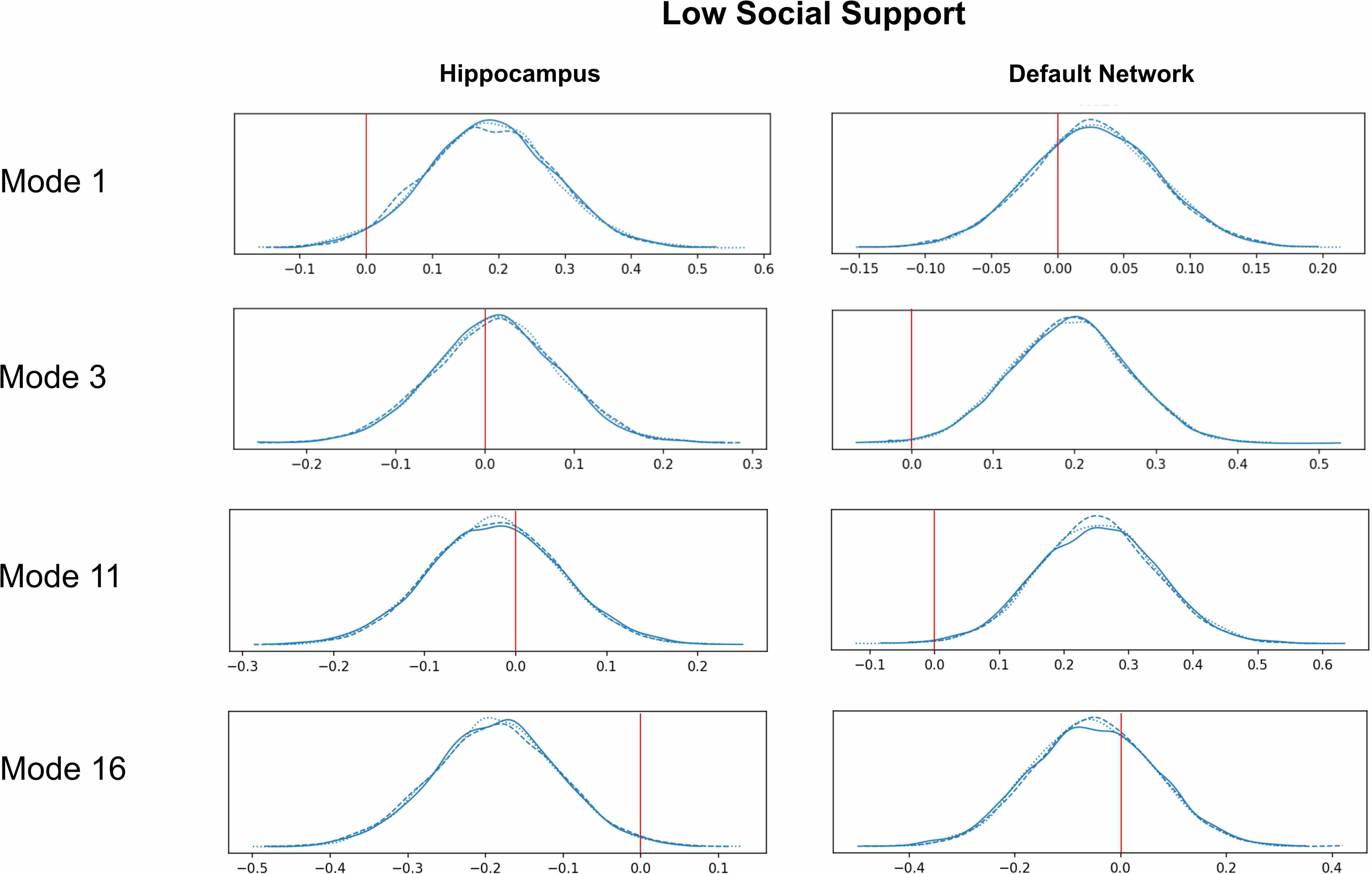

## References

Abdellaoui, A., Chen, H.Y., Willemsen, G., Ehli, E.A., Davies, G.E., Verweij, K.J., Nivard, M.G., de Geus, E.J., Boomsma, D.I., Cacioppo, J.T.J.J.o.p., 2019. Associations between loneliness and personality are mostly driven by a genetic association with neuroticism. 87, 386–397.

Alcalá-López, D., Smallwood, J., Jefferies, E., Van Overwalle, F., Vogeley, K., Mars, R.B., Turetsky, B.I., Laird, A.R., Fox, P.T., Eickhoff, S.B., 2018. Computing the social brain connectome across systems and states. Cerebral Cortex 28, 2207–2232.

Alexander, G.M., Farris, S., Pirone, J.R., Zheng, C., Colgin, L.L., Dudek, S.M., 2016. Social and novel contexts modify hippocampal CA2 representations of space. Nature communications 7, 1–14.

Alfaro-Almagro, F., Jenkinson, M., Bangerter, N.K., Andersson, J.L., Griffanti, L., Douaud, G., Sotiropoulos, S.N., Jbabdi, S., Hernandez-Fernandez, M., Vallee, E., 2018. Image processing and Quality Control for the first 10,000 brain imaging datasets from UK Biobank. Neuroimage 166, 400–424.

Ally, B.A., Hussey, E.P., Ko, P.C., Molitor, R.J., 2013. Pattern separation and pattern completion in Alzheimer’s disease: evidence of rapid forgetting in amnestic mild cognitive impairment. Hippocampus 23, 1246–1258.

Anacker, C., Luna, V.M., Stevens, G.S., Millette, A., Shores, R., Jimenez, J.C., Chen, B., Hen, R., 2018. Hippocampal neurogenesis confers stress resilience by inhibiting the ventral dentate gyrus. Nature 559, 98–102.

Andrews-Hanna, J.R., Reidler, J.S., Sepulcre, J., Poulin, R., Buckner, R.L., 2010. Functional-anatomic fractionation of the brain’s default network. Neuron 65, 550–562.

Andrews-Hanna, J.R., Smallwood, J., Spreng, R.N., 2014. The default network and self-generated thought: component processes, dynamic control, and clinical relevance. Annals of the New York Academy of Sciences 1316, 29.

Aronov, D., Nevers, R., Tank, D.W., 2017. Mapping of a non-spatial dimension by the hippocampal– entorhinal circuit. Nature 543, 719–722.

Atroszko, P., Pianka, L., Raczyńska, A., Sęktas, M., Atroszko, B., 2015. Validity and reliability of single-item self-report measures of social support.

Baker, S., Vieweg, P., Gao, F., Gilboa, A., Wolbers, T., Black, S.E., Rosenbaum, R.S., 2016. The human dentate gyrus plays a necessary role in discriminating new memories. Current Biology 26, 2629–2634.

Bakker, A., Kirwan, C.B., Miller, M., Stark, C.E., 2008. Pattern separation in the human hippocampal CA3 and dentate gyrus. Science 319, 1640–1642.

Behrens, T.E., Muller, T.H., Whittington, J.C., Mark, S., Baram, A.B., Stachenfeld, K.L., Kurth-Nelson, Z., 2018. What is a cognitive map? Organizing knowledge for flexible behavior. Neuron 100, 490–509.

Bellmund, J.L., Gärdenfors, P., Moser, E.I., Doeller, C.F., 2018. Navigating cognition: Spatial codes for human thinking. Science 362.

Berron, D., Schütze, H., Maass, A., Cardenas-Blanco, A., Kuijf, H.J., Kumaran, D., Düzel, E., 2016. Strong evidence for pattern separation in human dentate gyrus. Journal of Neuroscience 36, 7569–7579.

Biggio, F., Mostallino, M., Talani, G., Locci, V., Mostallino, R., Calandra, G., Sanna, E., Biggio, G., 2019. Social enrichment reverses the isolation-induced deficits of neuronal plasticity in the hippocampus of male rats. Neuropharmacology 151, 45–54.

Binder, J.R., Desai, R.H., Graves, W.W., Conant, L.L., 2009. Where is the semantic system? A critical review and meta-analysis of 120 functional neuroimaging studies. Cerebral Cortex 19, 2767–2796.

Boccara, C.N., Sargolini, F., Thoresen, V.H., Solstad, T., Witter, M.P., Moser, E.I., Moser, M.-B., 2010. Grid cells in pre-and parasubiculum. Nature neuroscience 13, 987–994.

Boomsma, D.I., Willemsen, G., Dolan, C.V., Hawkley, L.C., Cacioppo, J.T., 2005. Genetic and environmental contributions to loneliness in adults: The Netherlands Twin Register Study. Behavior genetics 35, 745–752.

Braga, R.M., Buckner, R.L., 2017. Parallel interdigitated distributed networks within the individual estimated by intrinsic functional connectivity. Neuron 95, 457–471. e455.

Braga, R.M., DiNicola, L.M., Becker, H.C., Buckner, R.L., 2020. Situating the left-lateralized language network in the broader organization of multiple specialized large-scale distributed networks. Journal of neurophysiology 124, 1415–1448.

Braga, R.M., Van Dijk, K.R., Polimeni, J.R., Eldaief, M.C., Buckner, R.L., 2019. Parallel distributed networks resolved at high resolution reveal close juxtaposition of distinct regions. Journal of neurophysiology 121, 1513–1534.

Buzsaki, G., 2006. Rhythms of the Brain. Oxford University Press.

Bzdok, D., Nichols, T.E., Smith, S.M., 2019. Towards algorithmic analytics for large-scale datasets. Nature Machine Intelligence 1, 296–306.

Chadwick, M.J., Bonnici, H.M., Maguire, E.A., 2014. CA3 size predicts the precision of memory recall. Proceedings of the National Academy of Sciences 111, 10720–10725.

Choi, S.W., Mak, T.S.-H., O'Reilly, P.F., 2020. Tutorial: a guide to performing polygenic risk score analyses. Nature Protocols 15, 2759–2772.

Cohen, S., Hoberman, H.M., 1983. Positive events and social supports as buffers of life change stress. Journal of Applied Social Psychology 13, 99–125.

Constantinescu, A.O., O'Reilly, J.X., Behrens, T.E., 2016. Organizing conceptual knowledge in humans with a gridlike code. Science 352, 1464–1468.

Courtney, A.L., Meyer, M.L., 2020. Self-other representation in the social brain reflects social connection. Journal of Neuroscience 40, 5616–5627.

Cyranowski, J.M., Zill, N., Bode, R., Butt, Z., Kelly, M.A.R., Pilkonis, P.A., Salsman, J.M., Cella, D., 2013. Assessing social support, companionship, and distress: National Institute of Health (NIH) Toolbox Adult Social Relationship Scales. Health psychology : official journal of the Division of Health Psychology, American Psychological Association 32, 293–301.

Danjo, T., Toyoizumi, T., Fujisawa, S., 2018. Spatial representations of self and other in the hippocampus. Science 359, 213–218.

Day, F.R., Ong, K.K., Perry, J.R.J.N.c., 2018. Elucidating the genetic basis of social interaction and isolation. 9, 1–6.

De Falco, E., Ison, M.J., Fried, I., Quiroga, R.Q., 2016. Long-term coding of personal and universal associations underlying the memory web in the human brain. Nature communications 7, 1–11.

DiNicola, L.M., Braga, R.M., Buckner, R.L., 2020. Parallel distributed networks dissociate episodic and social functions within the individual. Journal of neurophysiology 123, 1144–1179.

Dollinger, S.J., Malmquist, D., 2009. Reliability and validity of single-item self-reports: With special relevance to college students’ alcohol use, religiosity, study, and social life. Journal of General Psychology 136, 231–241.

Dranovsky, A., Picchini, A.M., Moadel, T., Sisti, A.C., Yamada, A., Kimura, S., Leonardo, E.D., Hen, R., 2011. Experience dictates stem cell fate in the adult hippocampus. Neuron 70, 908–923.

Dunbar, R.I., 1998. The social brain hypothesis. Evolutionary Anthropology: Issues, News, and Reviews: Issues, News, and Reviews 6, 178–190.

Dunbar, R.I., Shultz, S., 2007. Evolution in the social brain. Science 317, 1344–1347.

Efron, B., Tibshirani, R.J., 1994. An introduction to the bootstrap. CRC press.

Eichenbaum, H., Dudchenko, P., Wood, E., Shapiro, M., Tanila, H., 1999. The hippocampus, memory, and place cells: is it spatial memory or a memory space? Neuron 23, 209–226.

Elliott, J., Bodinier, B., Bond, T.A., Chadeau-Hyam, M., Evangelou, E., Moons, K.G., Dehghan, A., Muller, D.C., Elliott, P., Tzoulaki, I., 2020. Predictive accuracy of a polygenic risk score–enhanced prediction model vs a clinical risk score for coronary artery disease. Jama 323, 636–645.

Elliott, L.T., Sharp, K., Alfaro-Almagro, F., Shi, S., Miller, K.L., Douaud, G., Marchini, J., Smith, S.M., 2018. Genome-wide association studies of brain imaging phenotypes in UK Biobank. Nature 562, 210–216.

Fan, B.J., Bailey, J.C., Igo, R.P., Kang, J.H., Boumenna, T., Brilliant, M.H., Budenz, D.L., Fingert, J.H., Gaasterland, T., Gaasterland, D., 2019. Association of a primary open-angle glaucoma genetic risk score with earlier age at diagnosis. JAMA ophthalmology 137, 1190–1194.

Fanselow, M.S., Dong, H.-W., 2010. Are the dorsal and ventral hippocampus functionally distinct structures? Neuron 65, 7–19.

Frith, C.D., Frith, U., 2006. The neural basis of mentalizing. Neuron 50, 531–534.

Frith, U., Frith, C., 2010. The social brain: allowing humans to boldly go where no other species has been. Philos Trans R Soc Lond B Biol Sci 365, 165–176.

Gao, J., Davis, L.K., Hart, A.B., Sanchez-Roige, S., Han, L., Cacioppo, J.T., Palmer, A.A.J.N., 2017. Genome-wide association study of loneliness demonstrates a role for common variation. 42, 811–821.

Gauthier, J.L., Tank, D.W., 2018. A dedicated population for reward coding in the hippocampus. Neuron 99, 179–193. e177.

Gilbert, P.E., Kesner, R.P., Lee, I., 2001. Dissociating hippocampal subregions: A double dissociation between dentate gyrus and CA1. Hippocampus 11, 626–636.

Gould, E., McEwen, B.S., Tanapat, P., Galea, L.A., Fuchs, E., 1997. Neurogenesis in the dentate gyrus of the adult tree shrew is regulated by psychosocial stress and NMDA receptor activation. Journal of Neuroscience 17, 2492–2498.

Gould, E., Tanapat, P., McEwen, B.S., Flügge, G., Fuchs, E., 1998. Proliferation of granule cell precursors in the dentate gyrus of adult monkeys is diminished by stress. Proceedings of the National Academy of Sciences 95, 3168–3171.

Hafting, T., Fyhn, M., Molden, S., Moser, M.-B., Moser, E.I., 2005. Microstructure of a spatial map in the entorhinal cortex. Nature 436, 801–806.

Hartwigsen, G., Bengio, Y., Bzdok, D., 2021. How does hemispheric specialization contribute to human-defining cognition? Neuron.

Hawkley, L.C., Browne, M.W., Cacioppo, J.T., 2005. How Can I Connect With Thee?: Let Me Count the Ways. Psychological Science 16, 798–804.

Hitti, F.L., Siegelbaum, S.A., 2014. The hippocampal CA2 region is essential for social memory. Nature 508, 88–92.

HLA-C, H., 2009. Common polygenic variation contributes to risk of schizophrenia and bipolar disorder.

Ibi, D., Takuma, K., Koike, H., Mizoguchi, H., Tsuritani, K., Kuwahara, Y., Kamei, H., Nagai, T., Yoneda, Y., Nabeshima, T., 2008. Social isolation rearing-induced impairment of the hippocampal neurogenesis is associated with deficits in spatial memory and emotion-related behaviors in juvenile mice. Journal of neurochemistry 105, 921–932.

Iglesias, J.E., Augustinack, J.C., Nguyen, K., Player, C.M., Player, A., Wright, M., Roy, N., Frosch, M.P., McKee, A.C., Wald, L.L., 2015. A computational atlas of the hippocampal formation using ex vivo, ultra-high resolution MRI: application to adaptive segmentation of in vivo MRI. Neuroimage 115, 117–137.

Inouye, M., Abraham, G., Nelson, C.P., Wood, A.M., Sweeting, M.J., Dudbridge, F., Lai, F.Y., Kaptoge, S., Brozynska, M., Wang, T., 2018. Genomic risk prediction of coronary artery disease in 480,000 adults: implications for primary prevention. Journal of the American College of Cardiology 72, 1883–1893.

Jacobs, J., Weidemann, C.T., Miller, J.F., Solway, A., Burke, J.F., Wei, X.-X., Suthana, N., Sperling, M.R., Sharan, A.D., Fried, I., 2013. Direct recordings of grid-like neuronal activity in human spatial navigation. Nature neuroscience 16, 1188–1190.

Jamali, M., Grannan, B.L., Fedorenko, E., Saxe, R., Báez-Mendoza, R., Williams, Z.M., 2021. Single-neuronal predictions of others’ beliefs in humans. Nature, 1–5.

Jankowsky, J.L., Melnikova, T., Fadale, D.J., Xu, G.M., Slunt, H.H., Gonzales, V., Younkin, L.H., Younkin, S.G., Borchelt, D.R., Savonenko, A.V., 2005. Environmental enrichment mitigates cognitive deficits in a mouse model of Alzheimer’s disease. Journal of Neuroscience 25, 5217–5224.

Kanai, R., Bahrami, B., Roylance, R., Rees, G., 2012. Online social network size is reflected in human brain structure. Proceedings of the Royal Society B: Biological Sciences 279, 1327–1334.

Kesner, R.P., Rolls, E.T., 2015. A computational theory of hippocampal function, and tests of the theory: new developments. Neuroscience & Biobehavioral Reviews 48, 92–147.

Khera, A.V., Chaffin, M., Aragam, K.G., Haas, M.E., Roselli, C., Choi, S.H., Natarajan, P., Lander, E.S., Lubitz, S.A., Ellinor, P.T., 2018. Genome-wide polygenic scores for common diseases identify individuals with risk equivalent to monogenic mutations. Nature genetics 50, 1219–1224.

Knierim, J.J., Neunuebel, J.P., 2016. Tracking the flow of hippocampal computation: Pattern separation, pattern completion, and attractor dynamics. Neurobiology of learning and memory 129, 38–49.

Kogan, J.H., Franklandand, P.W., Silva, A.J., 2000. Long-term memory underlying hippocampus-dependent social recognition in mice. Hippocampus 10, 47–56.

Kraus, B.J., Brandon, M.P., Robinson II, R.J., Connerney, M.A., Hasselmo, M.E., Eichenbaum, H., 2015. During running in place, grid cells integrate elapsed time and distance run. Neuron 88, 578–589.

Krienen, F.M., Tu, P.-C., Buckner, R.L., 2010. Clan mentality: evidence that the medial prefrontal cortex responds to close others. Journal of Neuroscience 30, 13906–13915.

Kropff, E., Carmichael, J.E., Moser, M.-B., Moser, E.I., 2015. Speed cells in the medial entorhinal cortex. Nature 523, 419–424.

Kuchenbaecker, K.B., McGuffog, L., Barrowdale, D., Lee, A., Soucy, P., Healey, S., Dennis, J., Lush, M., Robson, M., Spurdle, A.B., 2017. Evaluation of polygenic risk scores for breast and ovarian cancer risk prediction in BRCA1 and BRCA2 mutation carriers. JNCI: Journal of the National Cancer Institute 109.

Laurita, A.C., DuPre, E., Ebner, N.C., Turner, G.R., Spreng, R.N., 2020. Default network interactivity during mentalizing about known others is modulated by age and social closeness. Social cognitive and affective neuroscience 15, 537–549.

Lecarpentier, J., Silvestri, V., Kuchenbaecker, K.B., Barrowdale, D., Dennis, J., McGuffog, L., Soucy, P., Leslie, G., Rizzolo, P., Navazio, A.S., 2017. Prediction of breast and prostate cancer risks in male BRCA1 and BRCA2 mutation carriers using polygenic risk scores. Journal of Clinical Oncology 35, 2240.

Leroy, F., Park, J., Asok, A., Brann, D.H., Meira, T., Boyle, L.M., Buss, E.W., Kandel, E.R., Siegelbaum, S.A., 2018. A circuit from hippocampal CA2 to lateral septum disinhibits social aggression. Nature 564, 213–218.

Leutgeb, J.K., Leutgeb, S., Moser, M.-B., Moser, E.I., 2007. Pattern separation in the dentate gyrus and CA3 of the hippocampus. Science 315, 961–966.

Leutgeb, S., Leutgeb, J.K., Treves, A., Moser, M.-B., Moser, E.I., 2004. Distinct ensemble codes in hippocampal areas CA3 and CA1. Science 305, 1295–1298.

Lever, C., Burton, S., Jeewajee, A., O'Keefe, J., Burgess, N., 2009. Boundary vector cells in the subiculum of the hippocampal formation. Journal of Neuroscience 29, 9771–9777.

Lewis, P.A., Rezaie, R., Brown, R., Roberts, N., Dunbar, R.I., 2011. Ventromedial prefrontal volume predicts understanding of others and social network size. Neuroimage 57, 1624–1629.

Li, S., Jin, M., Zhang, D., Yang, T., Koeglsperger, T., Fu, H., Selkoe, D.J., 2013. Environmental novelty activates β2-adrenergic signaling to prevent the impairment of hippocampal LTP by Aβ oligomers. Neuron 77, 929–941.

Liu, H., Stufflebeam, S.M., Sepulcre, J., Hedden, T., Buckner, R.L., 2009. Evidence from intrinsic activity that asymmetry of the human brain is controlled by multiple factors. Proceedings of the National Academy of Sciences 106, 20499–20503.

Lord, C., 2013. Aristotle’s” Politics”. University of Chicago Press.

MacDonald, C.J., Lepage, K.Q., Eden, U.T., Eichenbaum, H., 2011. Hippocampal “time cells” bridge the gap in memory for discontiguous events. Neuron 71, 737–749.

Mar, R.A., Mason, M.F., Litvack, A., 2012. How daydreaming relates to life satisfaction, loneliness, and social support: the importance of gender and daydream content. Consciousness and cognition 21, 401–407.

Marioni, R.E., Proust-Lima, C., Amieva, H., Brayne, C., Matthews, F.E., Dartigues, J.-F., Jacqmin-Gadda, H., 2015. Social activity, cognitive decline and dementia risk: a 20-year prospective cohort study. BMC public health 15, 1–8.

Mars, R.B., Neubert, F.-X., Noonan, M.P., Sallet, J., Toni, I., Rushworth, M.F., 2012. On the relationship between the “default mode network” and the “social brain”. Frontiers in human neuroscience 6, 189.

Mars, R.B., Sallet, J., Neubert, F.-X., Rushworth, M.F., 2013. Connectivity profiles reveal the relationship between brain areas for social cognition in human and monkey temporoparietal cortex. Proceedings of the National Academy of Sciences 110, 10806–10811.

Mashek, D., Cannaday, L.W., Tangney, J.P., 2007. Inclusion of community in self scale: A single-item pictorial measure of community connectedness. Journal of Community Psychology 35, 257–275.

McHugh, T.J., Jones, M.W., Quinn, J.J., Balthasar, N., Coppari, R., Elmquist, J.K., Lowell, B.B., Fanselow, M.S., Wilson, M.A., Tonegawa, S., 2007. Dentate gyrus NMDA receptors mediate rapid pattern separation in the hippocampal network. Science 317, 94–99.

Meira, T., Leroy, F., Buss, E.W., Oliva, A., Park, J., Siegelbaum, S.A., 2018. A hippocampal circuit linking dorsal CA2 to ventral CA1 critical for social memory dynamics. Nature communications 9, 1–14.

Meisner, A., Kundu, P., Zhang, Y.D., Lan, L.V., Kim, S., Ghandwani, D., Choudhury, P.P., Berndt, S.I., Freedman, N.D., Garcia-Closas, M., 2020. Combined Utility of 25 Disease and Risk Factor Polygenic Risk Scores for Stratifying Risk of All-Cause Mortality. The American Journal of Human Genetics 107, 418–431.

Mesoudi, A., Whiten, A., Dunbar, R., 2006. A bias for social information in human cultural transmission. British journal of psychology 97, 405–423.

Miller, K.L., Alfaro-Almagro, F., Bangerter, N.K., Thomas, D.L., Yacoub, E., Xu, J., Bartsch, A.J., Jbabdi, S., Sotiropoulos, S.N., Andersson, J.L., 2016. Multimodal population brain imaging in the UK Biobank prospective epidemiological study. Nat Neurosci 19, 1523.

Morton, N.W., Zippi, E.L., Noh, S., Preston, A.R., 2021. Semantic knowledge of famous people and places is represented in hippocampus and distinct cortical networks. Journal of Neuroscience.

Moser, E.I., Kropff, E., Moser, M.-B., 2008. Place cells, grid cells, and the brain’s spatial representation system. Annu. Rev. Neurosci. 31, 69–89.

Numssen, O., Bzdok, D., Hartwigsen, G., 2021. Functional specialization within the inferior parietal lobes across cognitive domains. Elife 10, e63591.

O’Keefe, J., Dostrovsky, J., 1971. The hippocampus as a spatial map: Preliminary evidence from unit activity in the freely-moving rat. Brain research.

O’keefe, J., Nadel, L., 1978. The hippocampus as a cognitive map. Oxford: Clarendon Press.

Oliva, A., Fernández-Ruiz, A., Leroy, F., Siegelbaum, S.A., 2020. Hippocampal CA2 sharp-wave ripples reactivate and promote social memory. Nature 587, 264–269.

Omer, D.B., Maimon, S.R., Las, L., Ulanovsky, N., 2018. Social place-cells in the bat hippocampus. Science 359, 218–224.

Parizkova, M., Lerch, O., Andel, R., Kalinova, J., Markova, H., Vyhnalek, M., Hort, J., Laczó, J., 2020. Spatial Pattern Separation in Early Alzheimer’s Disease. Journal of Alzheimer’s Disease, 1–18.

Penninkilampi, R., Casey, A.-N., Singh, M.F., Brodaty, H., 2018. The association between social engagement, loneliness, and risk of dementia: a systematic review and meta-analysis. Journal of Alzheimer’s Disease 66, 1619–1633.

Petrides, M., Tomaiuolo, F., Yeterian, E.H., Pandya, D.N., 2012. The prefrontal cortex: comparative architectonic organization in the human and the macaque monkey brains. cortex 48, 46–57.

Plachti, A., Eickhoff, S.B., Hoffstaedter, F., Patil, K.R., Laird, A.R., Fox, P.T., Amunts, K., Genon, S., 2019. Multimodal parcellations and extensive behavioral profiling tackling the hippocampus gradient. Cerebral Cortex 29, 4595–4612.

Quiroga, R.Q., Kraskov, A., Koch, C., Fried, I., 2009. Explicit encoding of multimodal percepts by single neurons in the human brain. Current Biology 19, 1308–1313.

Quiroga, R.Q., Reddy, L., Kreiman, G., Koch, C., Fried, I., 2005. Invariant visual representation by single neurons in the human brain. Nature 435, 1102–1107.

Radvansky, B.A., Dombeck, D.A., 2018. An olfactory virtual reality system for mice. Nature communications 9, 1–14.

Rechnitz, O., Slutsky, I., Morris, G., Derdikman, D., 2021. Hippocampal sub-networks exhibit distinct spatial representation deficits in Alzheimer’s disease model mice. Current Biology.

Rey, H.G., Gori, B., Chaure, F.J., Collavini, S., Blenkmann, A.O., Seoane, P., Seoane, E., Kochen, S., Quiroga, R.Q., 2020. Single neuron coding of identity in the human hippocampal formation. Current Biology 30, 1152–1159. e1153.

Sallet, J., Mars, R.B., Noonan, M., Andersson, J.L., O’reilly, J., Jbabdi, S., Croxson, P.L., Jenkinson, M., Miller, K.L., Rushworth, M.F., 2011. Social network size affects neural circuits in macaques. Science 334, 697–700.

Sarel, A., Finkelstein, A., Las, L., Ulanovsky, N., 2017. Vectorial representation of spatial goals in the hippocampus of bats. Science 355, 176–180.

Saxe, R., 2006. Uniquely human social cognition. Current opinion in neurobiology 16, 235–239.

Saxe, R., Kanwisher, N., 2003. People thinking about thinking people: the role of the temporo-parietal junction in “theory of mind”. Neuroimage 19, 1835–1842.

Schacter, D.L., Wagner, A.D., 1999. Medial temporal lobe activations in fMRI and PET studies of episodic encoding and retrieval. Hippocampus 9, 7–24.

Schaefer, A., Kong, R., Gordon, E.M., Laumann, T.O., Zuo, X.-N., Holmes, A.J., Eickhoff, S.B., Yeo, B.T., 2017. Local-global parcellation of the human cerebral cortex from intrinsic functional connectivity MRI. Cereb Cortex, 1–20.

Schurz, M., Radua, J., Aichhorn, M., Richlan, F., Perner, J., 2014. Fractionating theory of mind: a meta-analysis of functional brain imaging studies. Neuroscience & Biobehavioral Reviews 42, 9–34.

Schurz, M., Radua, J., Tholen, M.G., Maliske, L., Margulies, D.S., Mars, R.B., Sallet, J., Kanske, P., 2021a. Toward a hierarchical model of social cognition: A neuroimaging meta-analysis and integrative review of empathy and theory of mind. Psychological Bulletin 147, 293.

Schurz, M., Uddin, L., Kanske, P., Lamm, C., Sallet, J., Bernhardt, B., Mars, R., Bzdok, D., 2021b. Variability in brain structure and function reflects lack of peer support. Cerebral Cortex.

Scoville, W.B., Milner, B., 1957. Loss of recent memory after bilateral hippocampal lesions. Journal of neurology, neurosurgery, and psychiatry 20, 11.

Shen, L.-X., Yang, Y.-X., Kuo, K., Li, H.-Q., Chen, S.-D., Chen, K.-L., Dong, Q., Tan, L., Yu, J.-T., 2021. Social Isolation, Social Interaction, and Alzheimer’s Disease: A Mendelian Randomization Study. Journal of Alzheimer’s Disease, 1–8.

Silva-Gomez, A.B., Rojas, D., Juarez, I., Flores, G., 2003. Decreased dendritic spine density on prefrontal cortical and hippocampal pyramidal neurons in postweaning social isolation rats. Brain research 983, 128–136.

Smith, S.M., Douaud, G., Chen, W., Hanayik, T., Alfaro-Almagro, F., Sharp, K., Elliott, L.T., 2021. An expanded set of genome-wide association studies of brain imaging phenotypes in UK Biobank. Nature neuroscience, 1–9.

Spithoven, A., Cacioppo, S., Goossens, L., Cacioppo, J., 2019. Genetic contributions to loneliness and their relevance to the evolutionary theory of loneliness. Perspectives on Psychological Science 14, 376–396.

Spreng, R.N., Andrews-Hanna, J.R., 2015. The default network and social cognition. Brain mapping: An encyclopedic reference 1316, 165–169.

Spreng, R.N., Dimas, E., Mwilambwe-Tshilobo, L., Dagher, A., Koellinger, P., Nave, G., Ong, A., Kernbach, J.M., Wiecki, T.V., Ge, T., 2020. The default network of the human brain is associated with perceived social isolation. Nature communications 11, 1–11.

Stevenson, E.L., Caldwell, H.K., 2014. Lesions to the CA 2 region of the hippocampus impair social memory in mice. European Journal of Neuroscience 40, 3294–3301.

Stranahan, A.M., Khalil, D., Gould, E., 2006. Social isolation delays the positive effects of running on adult neurogenesis. Nature neuroscience 9, 526–533.

Strange, B.A., Witter, M.P., Lein, E.S., Moser, E.I., 2014. Functional organization of the hippocampal longitudinal axis. Nature Reviews Neuroscience 15, 655–669.

Taube, J.S., Muller, R.U., Ranck, J.B., 1990. Head-direction cells recorded from the postsubiculum in freely moving rats. I. Description and quantitative analysis. Journal of Neuroscience 10, 420–435.

Tavares, R.M., Mendelsohn, A., Grossman, Y., Williams, C.H., Shapiro, M., Trope, Y., Schiller, D., 2015. A map for social navigation in the human brain. Neuron 87, 231–243.

Tolman, E.C., 1948. Cognitive maps in rats and men. Psychological review 55, 189.

Vogel, J.W., La Joie, R., Grothe, M.J., Diaz-Papkovich, A., Doyle, A., Vachon-Presseau, E., Lepage, C., de Wael, R.V., Thomas, R.A., Iturria-Medina, Y., 2020. A molecular gradient along the longitudinal axis of the human hippocampus informs large-scale behavioral systems. Nature communications 11, 1–17.

Wang, H.-T., Smallwood, J., Mourao-Miranda, J., Xia, C.H., Satterthwaite, T.D., Bassett, D.S., Bzdok, D., 2020. Finding the needle in high-dimensional haystack: A tutorial on canonical correlation analysis. Neuroimage.

Watarai, A., Tao, K., Wang, M.-Y., Okuyama, T., 2021. Distinct functions of ventral CA1 and dorsal CA2 in social memory. Current opinion in neurobiology 68, 29–35.

Wesnes, K.A., Annas, P., Basun, H., Edgar, C., Blennow, K., 2014. Performance on a pattern separation task by Alzheimer’s patients shows possible links between disrupted dentate gyrus activity and apolipoprotein E∈ 4 status and cerebrospinal fluid amyloid-β42 levels. Alzheimer’s research & therapy 6, 1–8.

Wikenheiser, A.M., Gardner, M.P., Mueller, L.E., Schoenbaum, G., 2021. Spatial representations in rat orbitofrontal cortex. Journal of Neuroscience 41, 6933–6945.

Wisse, L.E., Daugherty, A.M., Olsen, R.K., Berron, D., Carr, V.A., Stark, C.E., Amaral, R.S., Amunts, K., Augustinack, J.C., Bender, A.R., 2017. A harmonized segmentation protocol for hippocampal and parahippocampal subregions: Why do we need one and what are the key goals? Hippocampus 27, 3–11.

Witten, D.M., Tibshirani, R., Hastie, T., 2009. A penalized matrix decomposition, with applications to sparse principal components and canonical correlation analysis. Biostatistics, kxp008.

Wood, E.R., Dudchenko, P.A., Robitsek, R.J., Eichenbaum, H., 2000. Hippocampal neurons encode information about different types of memory episodes occurring in the same location. Neuron 27, 623–633.

Wray, N.R., Lin, T., Austin, J., McGrath, J.J., Hickie, I.B., Murray, G.K., Visscher, P.M., 2021. From basic science to clinical application of polygenic risk scores: a primer. JAMA psychiatry 78, 101–109.

Xiang, X., Lai, P.H.L., Bao, L., Sun, Y., Chen, J., Dunkle, R.E., Maust, D., 2021. Dual Trajectories of Social Isolation and Dementia in Older Adults: A Population-Based Longitudinal Study. Journal of Aging and Health 33, 63–74.

Xu, T., Nenning, K.-H., Schwartz, E., Hong, S.-J., Vogelstein, J.T., Goulas, A., Fair, D.A., Schroeder, C.E., Margulies, D.S., Smallwood, J., 2020. Cross-species functional alignment reveals evolutionary hierarchy within the connectome. Neuroimage 223, 117346.

