## Supplementary Online Material for "Lacking social support is associated with structural divergences in hippocampus-default network co-variation patterns"

| **Mode** | **Explained Variance (rho * 100)** |
| --- | --- |
| 1 | 51.0 |
| 2 | 42.2 |
| 3 | 38.9 |
| 4 | 30.9 |
| 5 | 26.8 |
| 6 | 23.3 |
| 7 | 22.2 |
| 8 | 20.1 |
| 9 | 17.8 |
| 10 | 16.8 |
| 11 | 15.0 |
| 12 | 14.7 |
| 13 | 13.8 |
| 14 | 11.9 |
| 15 | 11.5 |
| 16 | 11.1 |
| 17 | 9.9 |
| 18 | 9.2 |
| 19 | 8.4 |
| 20 | 7.9 |
| 21 | 7.4 |
| 22 | 7.2 |
| 23 | 6.6 |
| 24 | 6.2 |
| 25 | 6.1 |

**Supplementary Table 1. Relative explained variance of each of our 25 candidate hippocampus-default network signatures.** The canonical correlation (as Pearson’s rho * 100) of each of our 25 candidate modes identified through a canonical correlation analysis of hippocampus and default network subregion volumes.

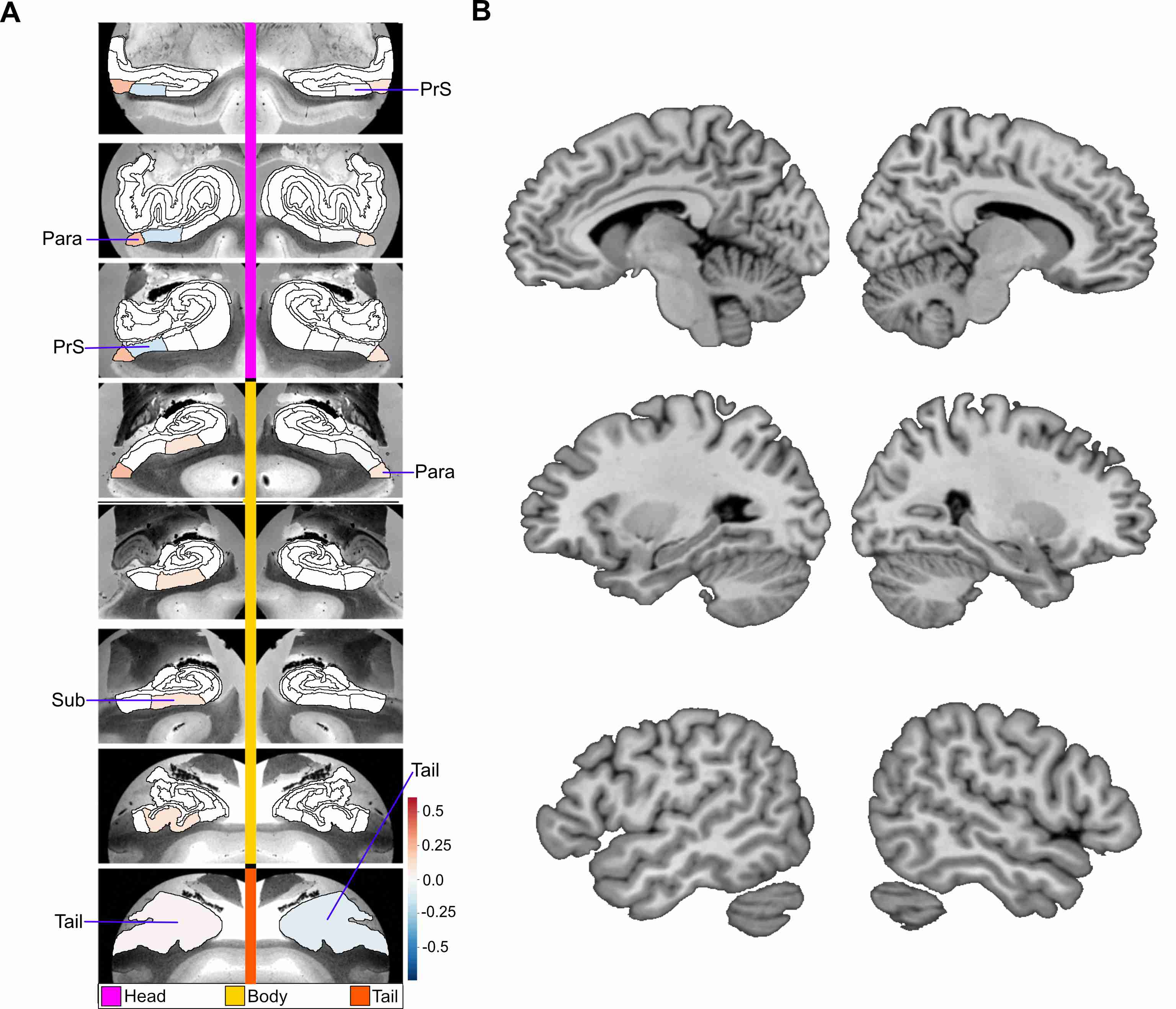

**Supplementary Figure 1. Low social support is associated with structural divergences preferentially in hippocampal tail and subicular subregions within mode 2 of hippocampus-default network co-variation.** Mode 2 of the CCA solution achieves the second most explanatory hippocampal-default network co-variation signature, with a canonical correlation of rho = 0.42. **A** shows the hippocampus (HC) subregion patterns (left, one canonical vector of mode 2) with parameter weights that robustly diverge between low and high social support groups; mapped onto 8 consecutive coronal slices of the left and right HC in the anterior (top) to posterior (bottom) direction. **B** shows the default network (DN) subregions patterns (right, other canonical vector of mode 2) that robustly diverge between the low and high social support groups. In the low social support group there are divergences in bilateral parasubiculum, bilateral presubiculum head, bilateral tail, and left subiculum body. However, in the DN there are no significant subregion divergences. These results demonstrate that in comparison to other HC-DN co-variation signatures, low social support primarily has effects on unique subregions within the second most explantory HC-DN signature. Para = parasubiculum, PrS = presubiculum, Sub = subiculum.

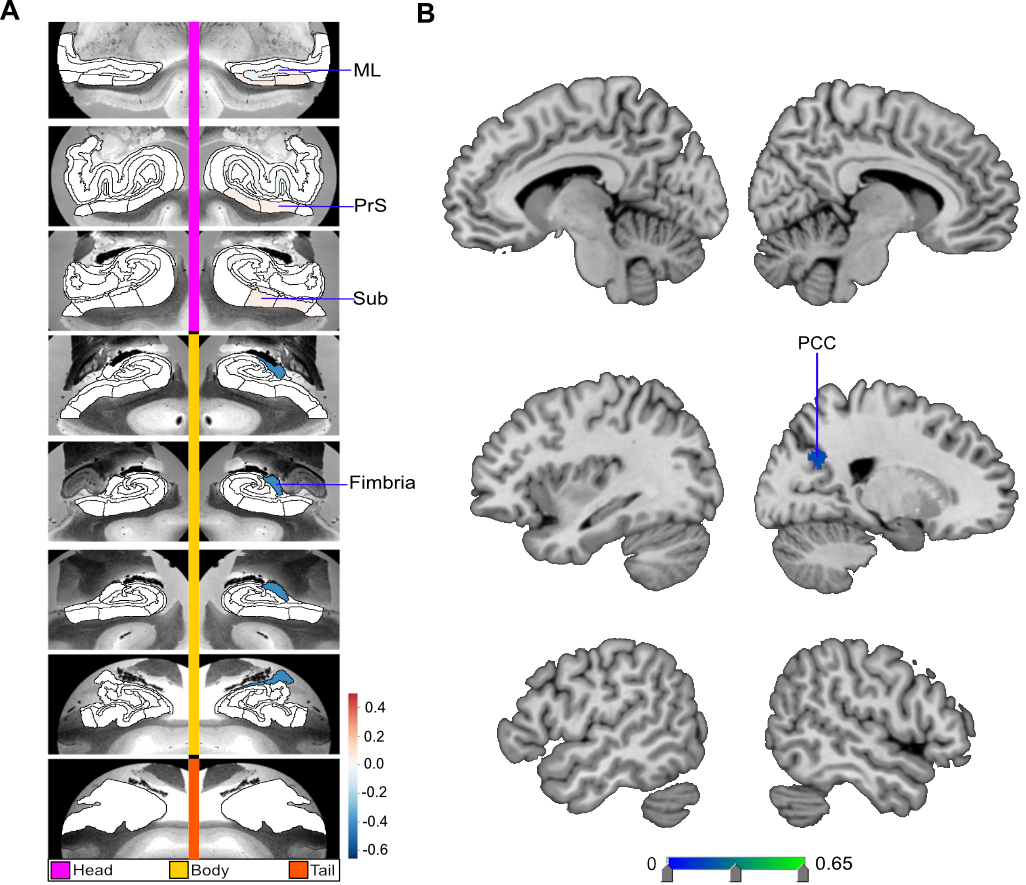

**Supplementary Figure 2. Low social support is associated with right hemispheric hippocampus and posterior cingulate divergences in mode 5.** Shown here are the subregion divergences in mode 5 of hippocampus-DN covariation. Mode 5 of the CCA solution acheives the fifth most explanatory hippocampal-default network co-variation, with a canonical correlation of rho = 0.27. **A** shows the hippocampus (HC) subregion patterns (left, one canonical vector of mode 5) with parameter weights that robustly diverge between low and high social support groups; mapped onto 8 consecutive coronal slices of the left and right HC in the anterior (top) to posterior (bottom) direction. **B** shows the default network (DN) subregions patterns (right, other canonical vector of mode 5) that robustly diverge between the low and high social support groups. Overall, in individuals with low social support there are HC divergences in right molecular layer head, right subiculum head, right presubiculum head, and right fimbria. There is also a DN subregion divergence in right posterior cingulate cortex (PCC). These results illustrate the selectivity of the alteration patterns within a particular mode. The results also demonstrate the tendency for lateralized divergences within modes. ML = molecular layer, PrS = presubiculum, Sub = subiculum.

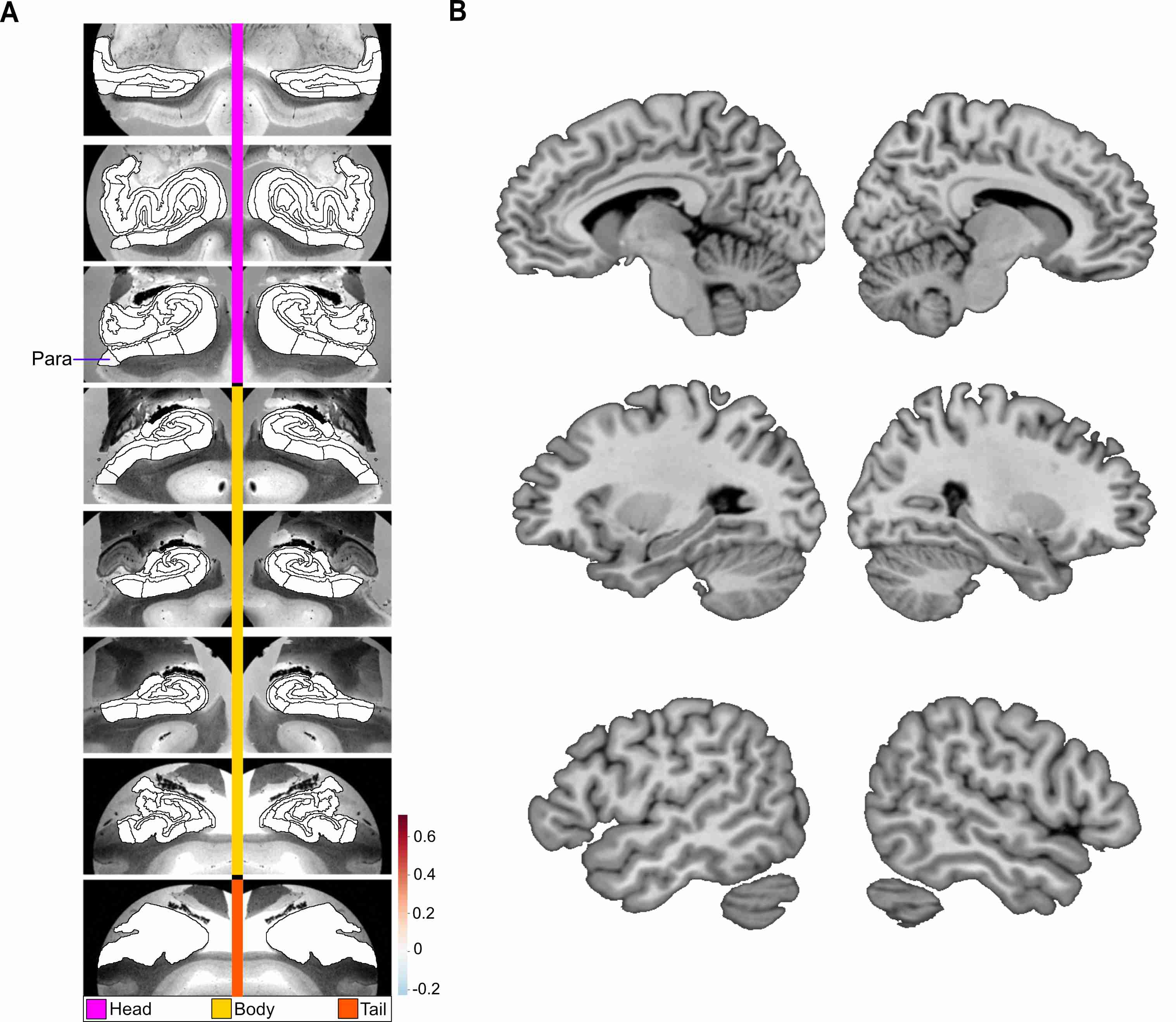

**Supplementary Figure 3. The parasubiculum alone shows significant structural divergence in a distinct signature of hippocampus-default network co-variation in individuals with low soccial support.** Mode 7 of the CCA solution achieves the seventh most explanatory hippocampal-default network co-variation signature, with a canonical correlation of rho = 0.22. **A** shows the hippocampus (HC) subregion patterns (left, one canonical vector of mode 7) with parameter weights that robustly diverge between low and high social support groups; mapped onto 8 consecutive coronal slices of the left and right HC in the anterior (top) to posterior (bottom) direction. **B** shows the default network (DN) subregions patterns (right, other canonical vector of mode 7) that robustly diverge between the low and high social support groups. Overall, the only divergence for the low social support group in this HC-DN signature was in the left parasubiculum (Para).
